# The structure of a Type III-A CRISPR-Cas effector complex reveals conserved and idiosyncratic contacts to target RNA and crRNA among Type III-A systems

**DOI:** 10.1101/2022.11.03.515080

**Authors:** Mohammadreza Paraan, Mohamed Nasef, Lucy Chou-Zheng, Sarah A. Khweis, Allyn J. Schoeffler, Asma Hatoum-Aslan, Scott M. Stagg, Jack A. Dunkle

## Abstract

Type III CRISPR-Cas systems employ multiprotein effector complexes bound to small CRISPR RNAs (crRNAs) to detect foreign RNA transcripts and elicit a complex immune response that leads to the destruction of invading RNA and DNA. Type III systems are among the most widespread in nature, and emerging interest in harnessing these systems for biotechnology applications highlights the need for detailed structural analyses of representatives from diverse organisms. We performed cryo-EM reconstructions of the Type III-A Cas10-Csm effector complex from *S. epidermidis* bound to an intact, cognate target RNA and identified two oligomeric states, a 276 kDa complex and a 318 kDa complex. 3.1 Å density for the well-ordered 276 kDa complex allowed construction of atomic models for the Csm2, Csm3, Csm4 and Csm5 subunits within the complex along with the crRNA and target RNA. We also collected small-angle X-ray scattering data which was consistent with the 276 kDa Cas10-Csm architecture we identified. Detailed comparisons between the *S. epidermidis* Cas10-Csm structure and the well-resolved bacterial (*S. thermophilus*) and archaeal (*T. onnurineus*) Cas10-Csm structures reveal differences in how the complexes interact with target RNA and crRNA which are likely to have functional ramifications. These structural comparisons shed light on the unique features of Type III-A systems from diverse organisms and will assist in improving biotechnologies derived from Type III-A effector complexes.

## Introduction

CRISPR-Cas systems provide adaptive immunity to prokaryotes by capturing fragments of genetic information from mobile genetic elements, such as plasmids and bacteriophage, and storing the information content in the spacers of a CRISPR array. The spacers are transcribed and processed into crRNAs, which when bound to effector Cas proteins, facilitate an interference response upon the detection of a complementary foreign nucleic acid [1, 2]. CRISPR-Cas systems have been organized into two classes, class 1 multi-Cas protein effectors versus class 2 single Cas protein effectors, and six types by bioinformatics analyses [3]. The *S. epidermidis* effector complex, known as Cas10-Csm, consists of a crRNA and five Cas proteins: Cas10, Csm2, Csm3, Csm4 and Csm5 (Fig. 1A). As a multi-protein effector complex, it is a member of class 1 and further categorized as Type III-A due to the presence of Cas10 and the small signature subunit Csm2.

**Figure 1.**
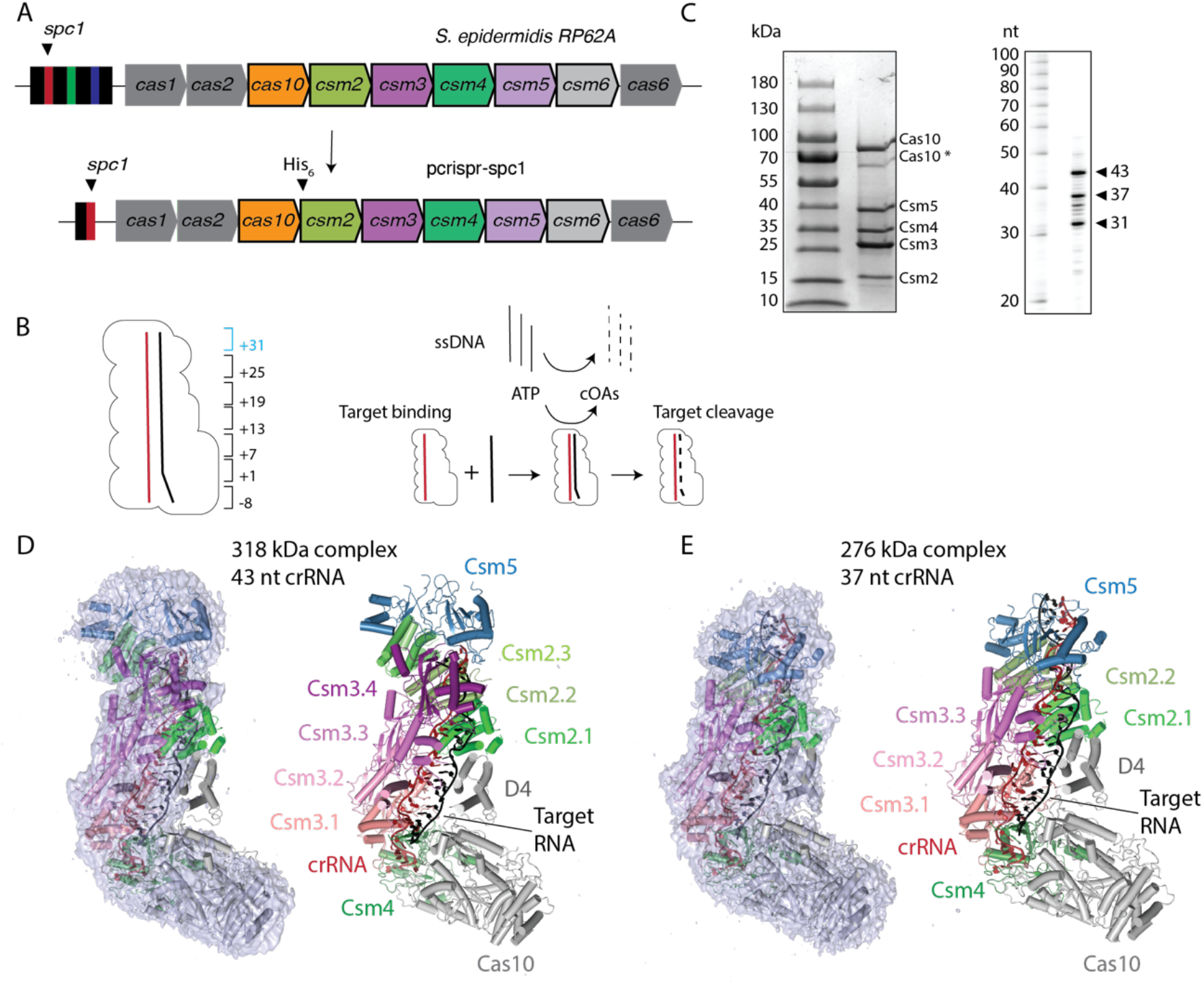
Overall architecture of two oligomeric states of *S. epidermidis* Cas10-Csm revealed by cryo-EM. (*A*) A schematic of the *S. epidermidis* Type III-A crispr locus. Cas10-Csm purification was facilitated by the construction of the plasmid *pcrispr-spc1*. (*B*) The numbering scheme describing positions within the crRNA-target duplex is shown. The six-nucleotide region beginning at +31 (blue) is present in the 43 nt cRNA complex but not the 37 nt crRNA complex. A schematic of steps in the interference reaction is shown. Target RNA binding activates single-stranded DNA (ssDNA) cleavage and cyclic oligoadenylates (cOAs) synthesis activities of Cas10-Csm. (*C*) SDS-PAGE and urea-PAGE analysis of purified Cas10-Csm. Cas10* denotes a band that has been identified by mass spectrometry as a truncated Cas10. CrRNA extracted from purified SeCas10-Csm possesses lengths of 31, 37 and 43 nt. (*D*) The larger complex consists of a 43 nt crRNA, three copies of Csm2 and four copies of Csm3. The cryo-EM density map is shown at left revealing that very little density is present for Cas10 domain 4 (D4) and Csm2.1. The map is contoured at level 3.0 σ. (*E*) The smaller complex consists of a 37 nt crRNA, two copies of Csm2 and three copies of Csm3. Density contoured at 3.0 σ is shown. In contrast to the larger complex, density is observed for Cas10 domain 4 and Csm2.1. A molecular model of the 276 kDa complex has been deposited with the PDB, code 8DO6.

Type III CRISPR-Cas systems are among the most abundant in nature and considered the most complex [3, 4]. Type III systems comprise approximately 25% of all CRISPR systems, a prevalance which suggests their physiological importance in prokaryotes [5]. There are six subtypes currently described (A-F), and among these, the III-A and III-B system are the best characterized [3]. These systems possess a distinct interference activity: upon sensing foreign RNA transcripts, the synthesis of second-messenger molecules, cyclic oligoadenylates (cOAs), is activated by the Palm-2 domain of Cas10 [6, 7]. The second-messenger binds to Csm6, a nuclease that is not a part of the Cas10-Csm complex, and activates its latent, indiscriminate RNase activity, which in turn drives the cell to dormancy to block viral replication (Fig. 1B) [6, 7]. Type III CRISPR systems also possess the ability to degrade foreign RNA complementary to the crRNA (target RNA) via the Csm3/Cmr4 protein, and many complexes also have the ability to degrade single-stranded (ss) DNA via the HD nuclease domain of Cas10, comprising a complex multifaceted interference response [8–10].

Recently, multiple investigators noted the intrinsic ability of Type III systems to specifically detect RNA and amplify this detection event by Cas10-mediated cOA synthesis makes them well suited to serve as a diagnostic tool for RNA viruses [11–14]. A Type III CRISPR complex was incubated with SARS-CoV-2 RNA initiating cOA synthesis. The cOA molelcules stimulated the RNase activity of Csm6 or a Csm6 homolog which then cleaved an fluorophore-quencher reporter RNA unleashing a fluorescent signal indicating the presence of the viral RNA [11–13]. Alternatively, NucC, a cOA stimulated DNase, was used in the place of Csm6 in a reaction that uses a fluorophore-quencher double-stranded DNA to report on the presence of viral RNA [14]. The Type III virus detection schemes achieved specificity similar to RT-qPCR assays but required coupling of an isothermal amplification step to achieve similar sensitivity, attomolar level [11–13]. The synthesis of cOA produces H^+^ and PP_i_ products during the course of the reaction and Santiago-Frangos and co-workers showed these molecules can also be sensed to report on the presence of viral RNA [11]. In all cases the Type III CRISPR based assays proceeded as rapid and isothermal reactions indicating a path to achieve sensitive and specific molecular diagnostics without expensive equipment - critical features for deployment in point-of-care settings. Realizing the full potential of this biotechnology will require a detailed understanding of the structure and mechanism of Type III CRISPR systems.

Structures of Type III CRISPR systems from several organisms of varying resolution are currently available. Low resolution Type III structures appeared in 2014, providing a good description of the location of the Cas proteins in the complex, and a high resolution structure of a Type III-B appeared shortly after [15–18]. Type III-A and Type III-B CRISPR systems differ structurally in two important ways: the presence of the small subunit protein Csm2 in Type III-A versus the presence of the small subunit protein Cmr5 in Type III-B and in the presence of the Cmr1 protein which produces a slightly different architecture adjacent to the 3’ end of the crRNA [5, 13, 16, 19]. Since the small subunit proteins, Csm2 and Cmr5, directly interact with target RNA upon its binding to the complex, it is likely these proteins play a critical role in sensing and activating interference [20, 21]. Therefore the first high resolution structures of Type III-A CRISPR systems in early 2019, were a welcome development [20, 22, 23]. These structures of Cas10-Csm from the archaebacterium *T. onnurineus* and the eubacterium *S. thermophilus* provided critical insights into the contacts of Cas10-Csm with crRNA and target RNA, but also differed in a key aspect: the *T. onnurineus* structure indicated little conformational change upon target binding, while the *S. thermophilus* structure underwent substantial conformational change upon target binding [20, 22]. In 2022, new high resolution structures of Cas10-Csm appeared. An *L. lactis* structure was reported further enabling structural biologists ability to compare and contrast across the available structures to formulate hypotheses regarding the mechanisms of target RNA sensing and activation [21]. While this manuscript was in preparation, high-resolution structures of *S. epidermidis* Cas10-Csm also were reported; however, in these structures, the cognate target RNA is not intact [24]. Thus, the confirmation of the complex in the active pre-cleavage state and corresponding contacts with the target RNA remain unclear.

To enrich the mechanistic understanding of how Type III systems function, we investigated the structure of the *S. epidermidis* Cas10-Csm complex bound to cognate target RNA by cryoelectron microscopy. A 3.1 Å map facilitated building atomic models for four of the five protein components, crRNA and target RNA. *S. epidermidis* Cas10-Csm is a leading model system in the study of Type III CRISPR systems dating to the earliest days of the field and therefore our structural model can assist in the design of detailed structure-function experiments utilizing the wealth of tools developed to study *S. epidermidis* Cas10-Csm [25, 26]. Additionally, we conducted a detailed comparison into the similarities and differences in how the *S. epidermidis* proteins interact with crRNA and target RNA compared to two well-resolved cognate, target RNA bound structures from *S. thermophilus* and *T. onnurineus*. Our observations shed light on the unique features of Type III-A systems from diverse organisms and are likely to guide efforts in optimizing the biotechnologies derived from Type III-A effector complexes

## Materials and Methods

### Modification of pcrispr to contain a single spacer

The *pcrispr-spc1* plasmid which contains a single repeat and *spc1* was constructed using a two-piece Gibson assembly with primers noted in Table S1. Briefly, the repeat-*spc1* region was amplified using primers F063/A010 from a plasmid containing the wild-type repeat-spacer array, and primers F062/L162 were used to amplify the backbone from a p*crispr-cas* plasmid lacking all repeats and spacers [27]. PCR products were then purified with EZNA Cycle Pure Kit (Omega Bio-tek) and Gibson assembled. The assembled construct was introduced into *S. aureus* RN4220 via electroporation, and transformants were subjected to PCR and DNA sequencing using primers A200/F111 to confirm the presence of a single repeat and spacer. The confirmed construct was then purified with EZNA Plasmid DNA Mini Kit (Omega Bio-tek) and introduced into *S. epidermidis* LM1680 strains via electroporation.

### Purification of the Cas10-Csm complex with bound crRNA

*S. epidermidis* LM1680 was transformed with *pcrispr-spc1* for expression of SeCas10-Csm complex. Cell growth, lysis and immobilized metal affinity chromatography (IMAC) were performed as described previously [28]. IMAC fractions were analyzed by SDS-PAGE and peak fractions were pooled, concentrated and loaded to a 5-20% w/v sucrose gradient for further purification by ultracentrifugation, a procedure previously reported [29]. Ultracentrifugation was performed for 41 hours in a SW32 rotor at 31,000 RPM and fractions were collected and analyzed by A280 and SDS-PAGE. Fractions with pure Cas10-Csm were pooled, concentrated to an A280 of 5.44, aliquoted and flash frozen in liquid nitrogen. Aliquots were stored at −80°C until needed.

### Structure determination of SeCas10-Csm by cryo-EM

Target RNA (ssRNA-01, Table S1) aliquots were thawed, heated at 70° for two minutes, then snap cooled on ice. SeCas10-Csm was thawed on ice, mixed with target RNA at a 1:1.6 molar ratio in the presence of 2 mM EDTA and incubated at 37°C for 5 minutes to form the target RNA bound SeCas10-Csm complex. The complex was then diluted 10-fold with 50 mM Tris-HCl pH 7.5, 150 mM NaCl buffer and applied to carbon coated grids for flash freezing. Movie frames were collected at the New York Structural Biology Center on a Titan Krios with a 300 keV FEG equipped with a Cs corrector, K3 camera, and a BioQuantum energy filter using Leginon at a pixel size of 0.846 Å with a total dose of 44.70 e/Å2 fractionated over 40 frames. About 5000 movie frames were collected over a period of 24 hours.

All image processing was performed in cryoSPARC 2 [30]. After patch motion correction and CTF estimation, the first set of particles were picked with the blob picker. A 2D classification was done on this set of particles and the resulting good 2D classes were used as templates for template matching. In order to remove false positives and bad particles, 2D classification followed by multiple rounds of ab initio and 3D heterogenous refinement were carried out. The strategy was to curate a set of good particles to be used for neural network training in Topaz [31]. After pooling together particles from different 3D classes, manual curation was done to create a set of 5,000 particles. This set of particles were used for Topaz training. Topaz cross validation was executed to optimize for the expected number of particles. The initial value for this parameter was set to 500 with 4 increments of the size 100. Training radius was set to 2. ResNet8 model architecture was trained for 15 epochs. After Topaz training, Topaz extract was carried out. With a particle threshold of −1, about 1.4 million particles were extracted and then binned by a factor of two. The rest of the processing pipeline is illustrated in supplementary figure S1. The dataset was heterogenous both in terms of stoichiometry and conformation. The smallest to the largest stoichiometry are labelled (complex I to III in figure S1). Complex II was used for the molecular models presented herein. Conformational heterogeneity was seen in Cas10 in 2D classifications. When refined to high resolution, complex II lost almost all the density for Cas10, but complex III retained the secondary structures. Variability analysis was used to tease out complex II and complex III [32]. This was confirmed by a heterogenous refinement. Final refinements were done using nonuniform refinement [33].

An atomic model the *S. epidermidis* Csm2-5 proteins with crRNA and target RNA was constructed using iterative manual modeling and real-space refinement in Coot followed by cycles of real-space refinement in Phenix [34, 35]. Docking of the coordinates for Csm3 and Csm2, described by PDB codes 6nbt and 6nbu, respectively, into the volume described in emd-27593 was performed in Coot. AlphaFold2 models of Csm4 and Csm5 (sequence references in Table S2) were remodeled within Coot to fit the volume [36, 37]. RNA molecules were built in Coot from the sequences described in Table S2. Molprobity was used to monitor coordinate geometry and minimize clashes [38].

### Collection of SEC-SAXS data

SEC-SAXS was performed at the SIBYLS beamline at the Advanced Light Source at Lawrence Berkeley National Laboratory. Three concentrations of SeCas10-Csm complex bound to target RNA (ssRNA-01) were formed by the brief incubation of SeCas10-Csm with a 1:1 mol:mol ratio of target RNA followed by application of the complex to a Shodex 804 SEC column flowing at 0.5 mL/min with running buffer of 50 mM Tris-HCl pH 7.5, 150 mM NH4Cl, 2% v/v glycerol. The concentrations of complex were 20 μM, 10 μM and 5 μM which corresponds to ~ 6 mg/mL, ~3 mg/mL and ~1.5 mg/mL. Frames collected at the SEC peak were averaged and buffer-subtracted using ScÅtter (https://bl1231.als.lbl.gov/scatter/). Subtracted curves were further analyzed using PRIMUS [39], and pairwise distance distribution functions were calculated using GNOM in PRIMUS [40]. Ab initio models were calculated using DAMMIN [41], and molecular models were calculated using SASREF [42]. To generate molecular models, coordinate files for individual subunits bound to RNA were generated by partitioning the *S. thermophilus* Cas10-Csm structure (PDB ID 6ifu) [20] into individual subunits bound to RNA segments and, in the case of the Csm1 subunit, partitioning the chain into two components at a domain boundary at position Lys-654 (Table S3). During rigid body model calculation, constraints were applied to limit distances between these components at covalent bonds linking nucleotides or the peptide backbone. CRYSOL [43] and OLIGOMER were used to generate theoretical scattering profiles from molecular models (those reported here and previously) and compare them to the experimental scattering profile. Because SAXS solutions are inherently degenerate and cannot distinguish enantiomorphs, mirror images of ab initio and molecular models are displayed wherever alignments with molecular models indicate that this is the biochemically relevant model. Enantiomorph transformation was performed using ALPRAXIN [44].

### Sequence logos and structural superpositions

The sequence logo describing Csm2 conservation was constructed using WebLogo [45]. Input was a multiple sequence alignment given by PFAM of all Uniprot entries in the protein family identified by PFAM, PF03750 [46]. The sequence logo for Csm5 was constructed using WebLogo from a multiple sequence alignment of 250 homologs to SeCsm5 identified by Protein BLAST and aligned with Clustal Omega. Structural superpositions were made in Pymol using the fit command with P atoms of the target or crRNA backbone selected.

## Results

### The overall architecture of *S. epidermidis* Cas10-Csm by cryo-EM and SAXS

The *S. epidermdis* Type III-A CRISPR array contains three spacer segments (*spc1-spc3*). To facilitate purification of an in vivo assembled Cas10-Csm complex with a single crRNA, the segments encoding *spc2* and *spc3* were removed from *pcrispr* (Fig. 1A). The *pcrispr* plasmid is designed for expression of *S. epidermidis* Cas10-Csm (SeCas10-Csm) in *S. epidermidis* cells and we have previously demonstrated this expression and purification approach leads to a ribonucleoprotein complex with robust activity in cyclic oligoadenylate synthesis and target RNA cleavage (Figs. 1B, S2) [29]. Cas10-Csm bound to a crRNA derived from *spc1* was purified and SDS-PAGE analysis demonstrated the presence of the five expected proteins. Urea-PAGE analysis of RNA extracted from the complex produced bands of the typically observed sizes of 31, 37 and 43 nucleotide-length crRNAs (Fig. 1C).

To obtain a cryo-EM structure of SeCas10-Csm bound to target RNA, purified complex was incubated with a slight excess of synthetic target RNA containing sequence for the nickase mRNA, which is complementary to *spc1* derived crRNA and is its natural target. SeCas10-Csm bound to the target was plunge frozen. Csm3 mediated target RNA cleavage was inhibitied during the incubation by the presence of EDTA, which chelates divalent metal ions required for the cleavage reaction. From 5,000 micrographs, 1.4 million particles were extracted and through the classificaion and refinement steps detailed in Figure S1, two distinct volumes at 3.1 Å resolution were reconstructed. Docking of the *S. epidermidis* Csm2 and Csm3 crystal structures, given by PDB codes 6nbu and 6nbt respectively, and manual building of crRNA, target RNA, Csm4 and Csm5 into the two volumes revealed that each corresponded to a different oligomeric state of SeCas10-Csm, a 318 kDa complex and a 276 kDa complex (Figs. 1D, 1E). The larger complex is composed of a 43 nt crRNA and an extra copy of Csm2 and Csm3 compared to the 276 kDa complex which possesses a 37 nt crRNA (Figs. 1D, 1E). An additional difference between the complexes is present: Csm2 is well ordered in the 276 kDa complex but, while present, poorly ordered in the 318 kDa complex (Figs. 1D, 1E). Cas10 is present in both complexes, but is highly flexible with a hinge motion observed in 2D classes. The flexibility of Cas10 led to density that is not of sufficient quality to support the construction of an atomic model for the protein. Since the 276 kDa complex possesses high quality density for crRNA, target RNA and proteins Csm2-5 an atomic model for these components was constructed (Fig. S3, Table S4). The complex was completed by docking of a homology model of the SeCas10 component for which there is agreement on the underlying secondary structures and domain positions (Fig. S4). A C_α_ model of the 318 kDa complex was constructured by docking the molecular models of the Csm2-5 coordinates and the Cas10 homology model into density. The molecular model of the 276 kDa complex faciliates identifying the detailed interactions of the Csm2, Csm3, Csm4 and Csm5 proteins with crRNA and target.

We performed a complementary analysis of the SeCas10-Csm architecture utilizing size-exclusion chromatography coupled with small-angle x-ray scattering (SEC-SAXS). Purified SeCas10-Csm bearing a single *spc1*-derived crRNA was purified and incubated with target RNA in a manner nearly identical to our cryo-EM experiments, then passed over a SEC column and subjected to in-line SAXS data collection. Data frames from the center of the SEC peak were used to generate a scaled, averaged scattering curve. This scattering curve was then used to generate sets of independently calculated *ab initio* envelopes and rigid body models of the SeCas10-Csm complex based on the previously published *S. thermophilus* 6ifu structure [20]. Replicate rigid body models showed good agreement with each other and with the set of *ab initio* envelopes (Figs. 2A, 2B, S5, S6; Table S5). We further compared our SAXS data and models to both the 276 kDa and 318 kDa EM models reported here. We found that our SAXS-derived *ab initio* and rigid body models both showed better agreement with the 276 kDa complex than the 318 kDa complex (Fig. 2C, Table S6). To further validate the agreement between the solution scattering data and the 276 kDa complex, we calculated theoretical SAXS profiles (via CRYSOL) from the 276 kDa and 318 kDa EM structures and compared them to our experimental SAXS data. The profile calculated from the 276 kDa model showed significantly better agreement with our experimental SAXS data than did the 318 kDa model profile (Fig. 2D, Table S7). Finally, attempts to deconvolute the experimental scattering curve (via OLIGOMER) with multiple theoretical scattering profiles generated from plausible molecular models consistently revealed the presence of only a single component with the same stoichiometry as the 276 kDa complex (Fig. S7). The low resolution afforded by SAXS reconstructions (as well as the inherent degeneracy of three-dimensional structural solutions derived from two-dimensional scattering data) do not allow us to unambiguously confirm the stoichiometry of the complex using SAXS data alone, but the agreement between the 276 kDa complex and our SAXS-derived models suggests that the 276 kDa model stoichiometry exists and may predominate in solution.

**Figure 2.**
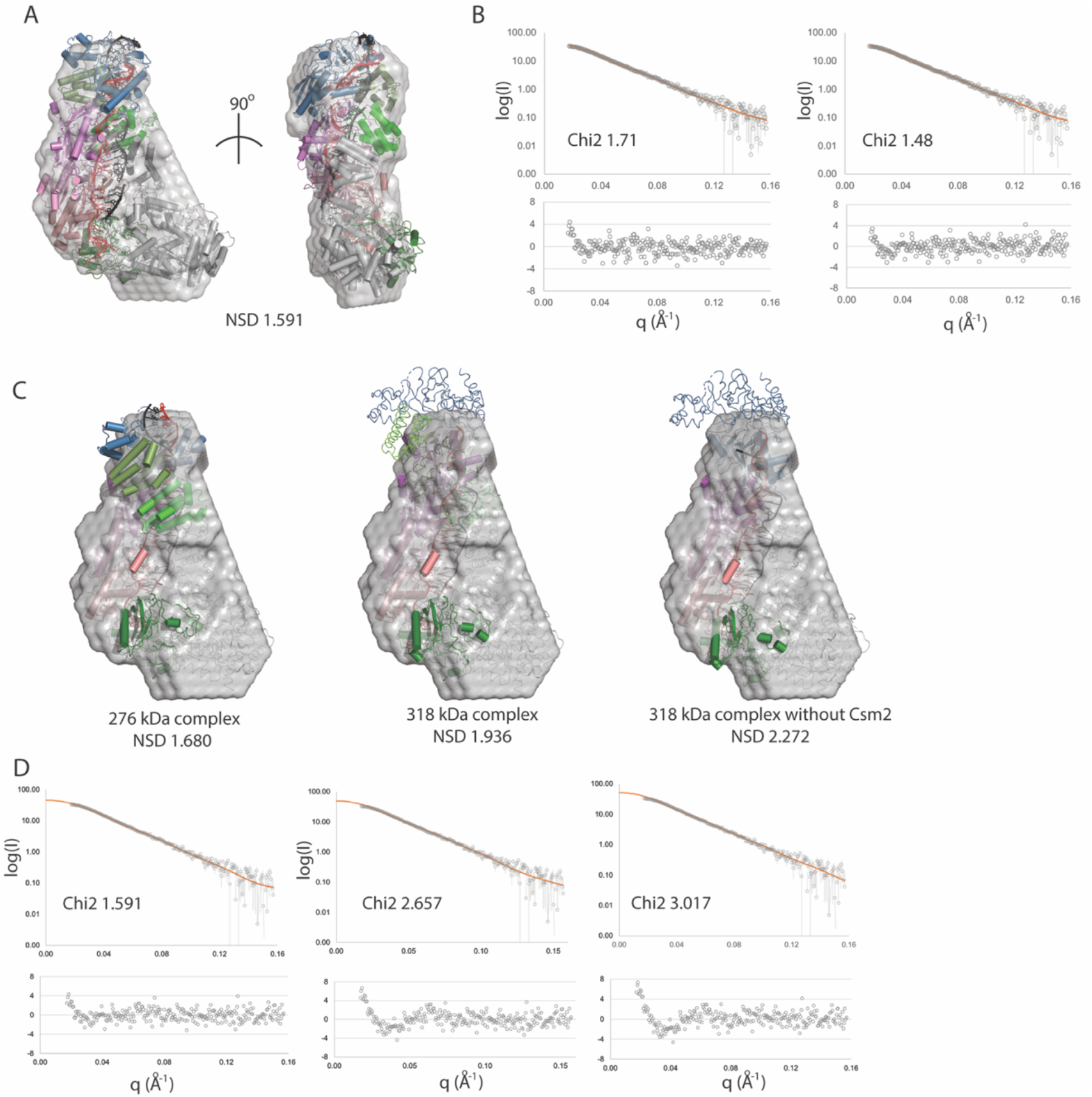
Support for Cas10-Csm complex stoichiometry from SEC-SAXS. (*A*) Multiple *ab initio* and rigid body models were calculated from the averaged peak frames of the SEC-SAXS run. Representative *ab initio* (gray envelope) and rigid body (cartoon) models are shown superposed (via SASREF). (*B*), The fits of the models from panel *A* (orange) to the experimental scattering curve (gray) are shown along with error-weighted residuals (lower panels). The rigid body model fit is shown at left, the *ab initio* model, at right. (*C*) EM structures reported here, overlaid with a representative *ab initio* model from our SAXS analysis. (*D*) CRYSOL was used to generate theoretical scattering curves from the EM models reported in this study, and these theoretical curves (orange) were scaled to the experimental SAXS data (gray). From left to right, the models shown are the 276 kDa complex, the 318 kDa complex, and the 318 kDa complex minus Csm2 subunits. As in *B*, error-weighted residuals are shown below each plot. Throughout this figure normalized spatial discrepancy is abbreviated NSD.

Superposition of the coordinates of the 276 kDa complex and the 318 kDa derived from cryo-EM reveal nearly identical positioning of Csm4 and the Csm3 oligomer. The Csm2 oligomer shifts slightly toward Cas10 in the 276 kDa complex a movement that could indicate sampling of a different conformational state, the influence of the differing stoichiometry or a combination of both (Fig. S8A). A superposition of the SAXS-derived rigid-body model that best fit the ab initio envelope to the 276 kDa complex also reveals that the 276 kDa complex has a slight shift of the Csm2 oligomer towards Cas10 (Fig. S8B). The conformational differences observed are not surprising given the documented dynamic nature of the Cas10-Csm complex [20, 21, 47]. However, a further detailed analysis of the differences cannot be conducted at this time due to the entanglement of stoichiometric differences and biases arising from the differing techniques used, SAXS versus cryo-EM.

### The detailed interactions of the Csm2-5 proteins with crRNA and target RNA

We used the atomic models of the Csm2, Csm3, Csm4 and Csm5 proteins to answer two questions. First, in *S. epidermidis* Cas10-Csm what are the specific interactions each of these proteins make with target and crRNA? Second, to what degree are the interactions between the Csm proteins and target and crRNA conserved versus idiosyncratic among Cas10-Csm complexes? To address this question we used structures available for another bacterial Cas10-Csm (*S. thermophilus*, PDB code 6ifu) and archaeal Cas10-Csm (*T. onnurineus*, PDB code 6mus). The second question is important because it directly bears on the larger question: to what degree is there a conserved mechanism for sensing the binding of target to crRNA and activating interference in Type III-A CRISPR-Cas? Recent biochemical data has identified site-directed mutants of Csm2, Csm3 and Csm4 with specific defects in interference and this literature will be described below as necessary [20–22].

Csm2 is present in two copies in the 276 kDa complex, contacting target RNA in the vicinity of positions +13 to +24 (Figs. 1B, 3A). Helix-α2 is positioned in the major groove of the target-crRNA duplex and conserved residues within the N-terminal region of α2 participate in hydrogen bonding and electrostatic interactions with the sugar-phosphate backbone of target RNA (Figs. 3B, 3C). Two conserved residues in α3, Y87 and R91, interact with phosphates at the +23 and +22 positions of target RNA, respectively (Csm2.2 protomer, Fig. 3C). Superposition of the bacterial Cas10-Csm complex from *S. thermophilus* with the *S. epidermidis* complex shows nearly identical interactions between Csm2 and target (Fig. 3C). A superposition of *T. onnurineus* Cas10-Csm with the *S. epidermidis* complex, however, reveals substantial differences in how Csm2 contacts target RNA (Fig. 3D). ToCsm2 residues Y129 and K133 in α3 are equivalent to SeCsm2 residues Y87 and R91 yet the ToCsm2 residues are located greater than 8 Å away from target RNA (Fig. 3D). This is due to a substantial difference in how Csm2 is positioned within the ToCas10-Csm complex that also affects ToCsm2 α2: the N-terminal residues of α2 contact target RNA but contact the +18 position rather than the +22 and +23 positions and make fewer overall contacts to target RNA (Fig. 3D). The Arg residue equivalent to Csm2 R49 (ToCsm2 R96) has been analyzed by site-directed mutagenesis in *S. thermophilus* Cas10-Csm and *L. lactis* Cas10-Csm and shown to be critical for target RNA cleavage activity confirming that the Csm2 contacts to target RNA have functional importance [20, 21].

**Figure 3.**
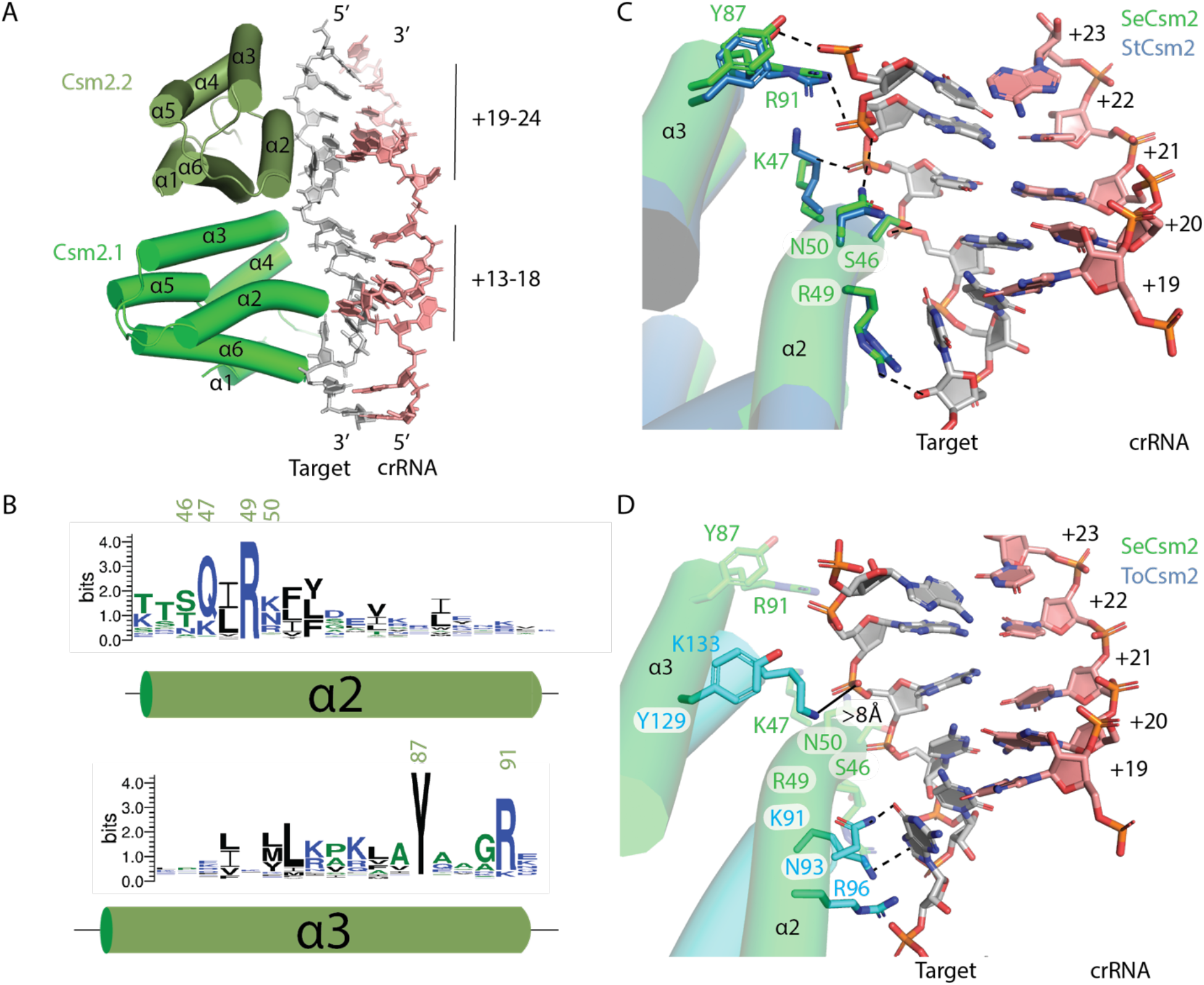
Conserved and idiosyncratic interactions of Csm2 with target RNA. (*A*) Helices α2 and α3 of *S. epidermidis* Csm2 (SeCsm2) form extensive interactions with target RNA in the major groove of the target-crRNA duplex. (*B*) A sequence logo depicting the conservation of α2 and α3 of Csm2. Residues of interest are numbered according to the SeCsm2 sequence. (*C*) A superposition of *S. thermophilus* Csm2 (StCsm2), given by PDB code 6ifu, with SeCsm2 was performed using the phosphate atoms of six nucleotides of target RNA (+19 to +24 positions). The superposition reveals that α2 and α3 in SeCsm2 and StCsm2 interact with target RNA in a similar manner. The target RNA-crRNA duplex depicted is from the SeCas10-Csm structure. (*D*) A superposition of *T. onnurineus* Csm2 (ToCsm2), given by PDB code 6MUS, with SeCsm2 was performed as in (*C*) which reveals that α2 and α3 in ToCsm2 are translated in space relative to SeCsm2, form fewer interactions with target RNA and these differ from the pattern seen in StCsm2 and SeCsm2. K91 of ToCsm2 forms a cation-pi interaction with the target-RNA nucleotide at position +18, analogous to the interaction of SeCsm2 R49 with the +18 position.

Csm3, like other Cas7 family proteins, forms a filament that cradles crRNA presenting it to solvent (Fig. 1E). Csm3 is composed of an RNA recognition motif (RRM) core elaborated with an RNase loop following β1, the α2 region and the thumb (or hook) region (Figs. 4A, 4B). The α2 loop contacts the minor groove of the crRNA-target duplex and the RNase loop contacts the major groove. The RNase loop performs divalent metal dependent target RNA cleavage (Figs. 4A, 4B). The thumb region inserts into the duplex disrupting nucleotide stacking (Figs. 4A, 4B). The bacterial Csm3 proteins, SeCsm3 and StCsm3, are very similar (Table S8). However, the archaeal, ToCsm3, possesses a more extensive α2 region (Fig. 4C).

**Figure 4.**
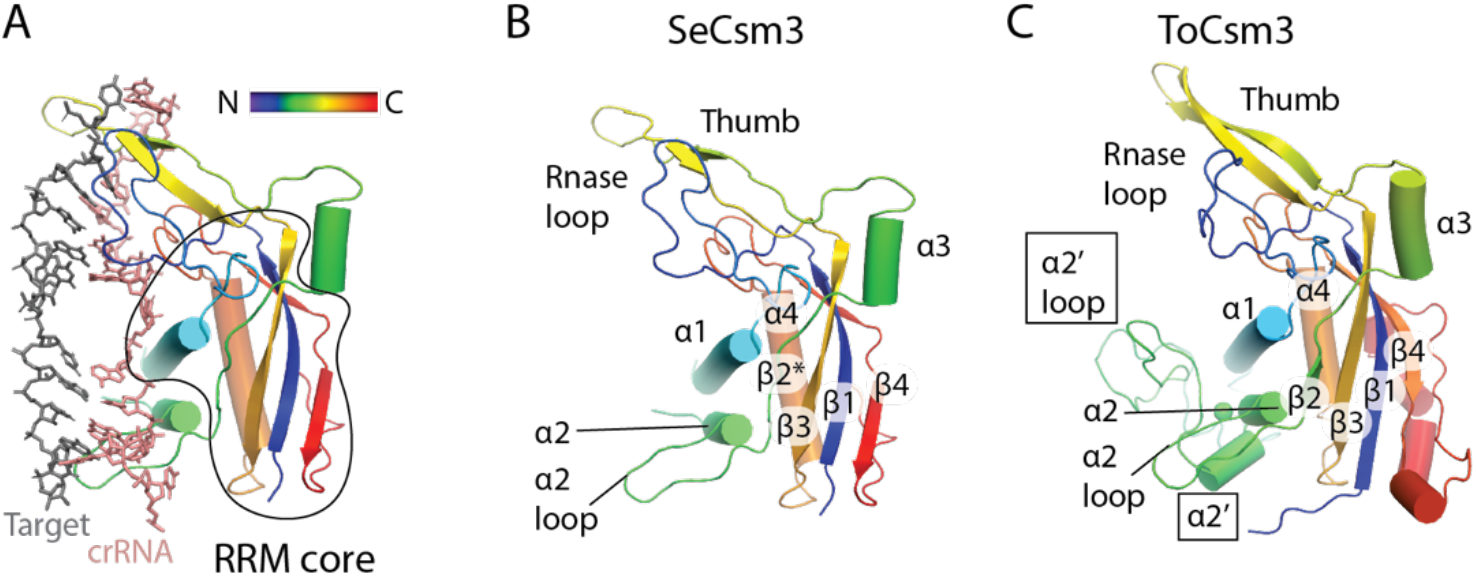
Variation in the secondary structure of Csm3. (*A*) The hydrophobic core of Csm3 is composed of an RNA recognition motif (RRM core). The secondary structure of Csm3 is colored by the position of each element on the chain from the N-terminus (blue) to the C-terminus (red). (*B*) The secondary structure of *S. epidermdis* Csm3 (SeCsm3) is shown with key functional components highlighted: the RNase loop contains Asp 32 which coordinates a metal to catalyze target RNA cleavage, α1 forms key interactions with crRNA, the α2 loop interacts with crRNA and lastly the thumb, composed of a β-hairpin protrudes through the crRNA-target RNA duplex interrupting it. β2* indicates that SeCsm3 adopts only β-strand like geometry at the position of β2 in the canonical RRM fold. (*C*) The secondary structure of *T. onnurineus* Csm3 (ToCsm3) differs from SeCsm3 because it contains additional elements of interest including the α2’ helix and the α2’ loop which make contacts to the target RNA that do not occur in SeCsm3. The structure depicted is taken from PDB code 6mus.

The SeCas10-Csm 276 kDa complex possesses three copies of Csm3 which have similar interactions with crRNA. Our analysis below focuses on Csm3.3 contacts to crRNA. The majority of contacts to crRNA occur via α1 residues. K52 and R56 make electrostatic interactions with the backbone of crRNA while S49, K54 and N57 contact the flipped nucleotide (+12 for Csm3.3, Fig. 5A). Thumb residues, N125 and I127 make additional contacts to the +12 crRNA position (Fig. 5A). The α2 region can make a single hydrogen bond to the minor groove via S86. The D32 residue of the RNase loop, which has been shown to be critical for target cleavage, is visible in the structure but is not coordinating a divalent metal ion, as it is expected to, because of the EDTA treatment of our sample prior to structural analysis (Figs. 5A, 5B).

**Figure 5.**
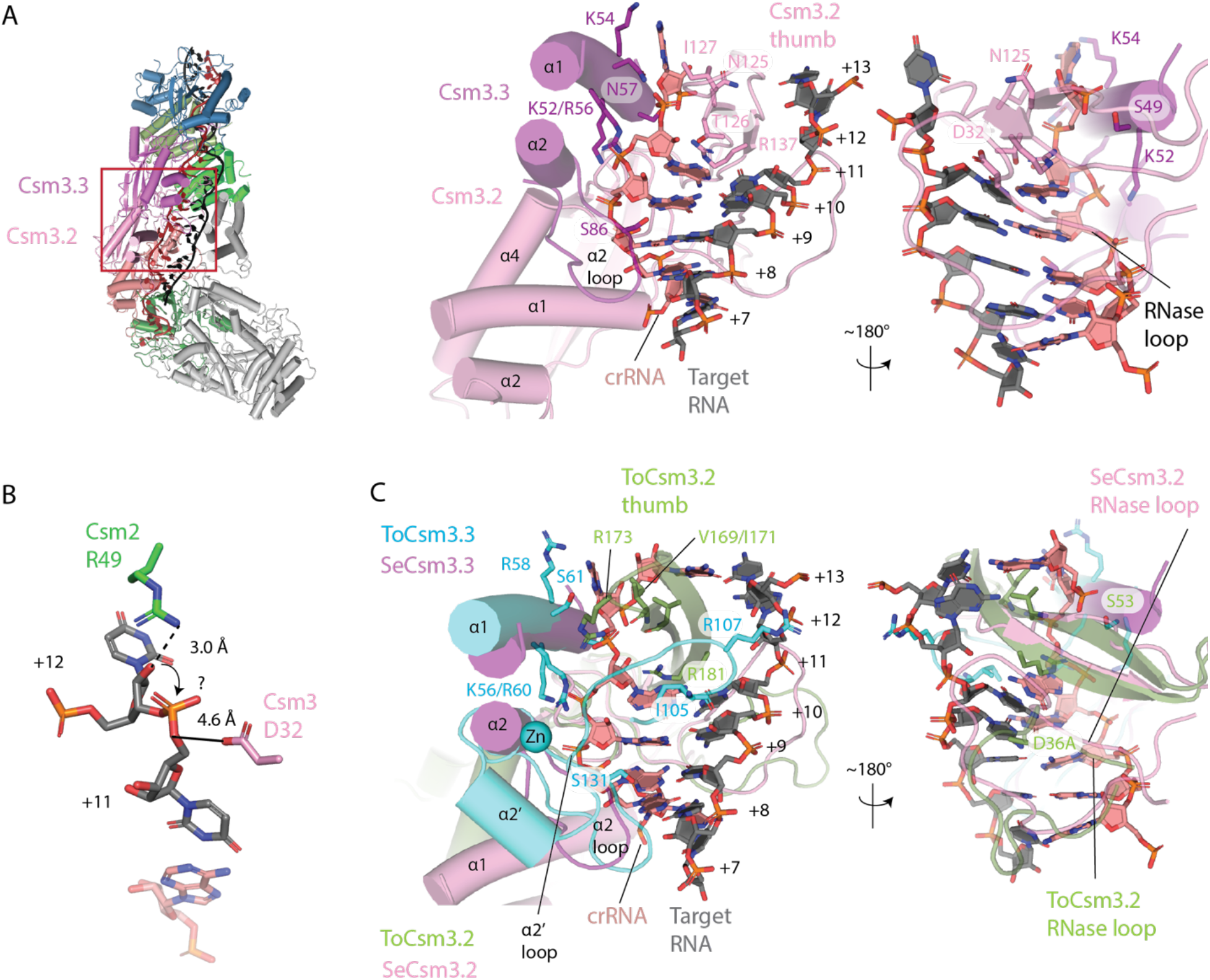
*S. epidermidis* Csm3 only contacts crRNA, however, *T. onnurineus* Csm3 contacts both crRNA and target RNA, utilizing the α2’ loop. (*A*) A red rectangle marks the area of Cas10-Csm which is magnified to the right. The detailed interactions of *S. epidermidis* Csm3 with crRNA are shown. A basic patch in α-helix 1 interacts with the terminal paired and the unpaired crRNA nucleotides of the repeating hexa-nucleotide structure. The S86 residue in the α2 loop hydrogen bonds to the minor groove of the duplex and residues N125-I127 of the Csm3.2 thumb form van der Waal’s interactions and hydrogen bonds with crRNA. An extensive loop structure that follows β-strand 1 of Csm3 and sits on the major groove of the crRNA-target duplex, is termed the RNase loop because it contains the catalytically essential residue D32. (*B*) Csm2 R49 and Csm3 D32 are positioned to catalyze target RNA cleavage but the mechanism of cleavage, in-line attack (arrow) or hydrolysis is unclear. (*C*) A superposition of *T. onnurineus* Csm2 (ToCsm3), given by PDB code 6mus, is shown revealing similar interactions of α-helix 1 and the α2 loop with crRNA in the two structures. However, ToCsm3 uses the α2’ loop, a structure not present in SeCsm2, to form hydrogen bonds (I105) and electrostatic interactions (R107) to target RNA. Additionally, the ToCsm3 thumb interacts differently with crRNA utilizing an electrostatic interaction via R173 that is absent in SeCsm3. The crRNA-target duplex shown is from PDB code 6mus.

A comparison of bacterial to archaeal Csm3, reveals that ToCsm3 has similar interactions to crRNA along α1, however, ToCsm3 uses an electrostatic interaction from thumb residue R173 to stabilize the +12 position flipped nucleotide (Fig. 5B). A substantial difference is seen in the addition of the α2’ region in ToCsm3 (Figs. 4C, 5B). The α2’ loop spans the minor groove of crRNA-target duplex and contacts target RNA via residues I105 and R107 (Fig. 5B). Since the bacterial Csm3 structures do not possess the α2’ region and contact only crRNA, not target, the ToCsm3 α2’ region could be the source of a substantial mechanistic difference between archaeal and bacterial Cas10-Csm [20, 22].

Csm4 is a Cas5 family protein, which in both Type I and Type III CRISPR systems, interacts with the 5’ end of crRNA [3, 48]. Like Csm3, Csm4 consists of an RRM core with a thumb region. Helices α3 and α4 are present in both SeCsm4 and ToCsm4, but interact differently with Cas10 in each case (Figs. 6A, 6B). The 5’ nt of crRNA, position −8 is gripped by SeCsm4 via pi-stacking with residues F40 and H291 (Figs. 1B, 7A). The −7 nt is splayed across α1 with a hydrogen-bonding interaction between Q251 and R191 contributing to nt positioning by steric occlusion (Fig. 7A). F249 stabilizes an A-form helix segment that spans the −5 to −2 positions of crRNA by pi-stacking to the −5 nt while Y148 pi-stacks to the −2 nt (Fig. 7A). Van der Waal’s interactions of residues H17, L23 and R190 also appear to contribute to formation of the A-form geometry of the −5 to −2 nts, a geometry that is pre-formed to inspect the 3’ flank region of target RNA for complementarity [20, 22, 49]. Residues Y148, V132 and L134 appear to promote the flipped conformation of the −1 nt by steric occlusion (Fig. 7A).

**Figure 6.**
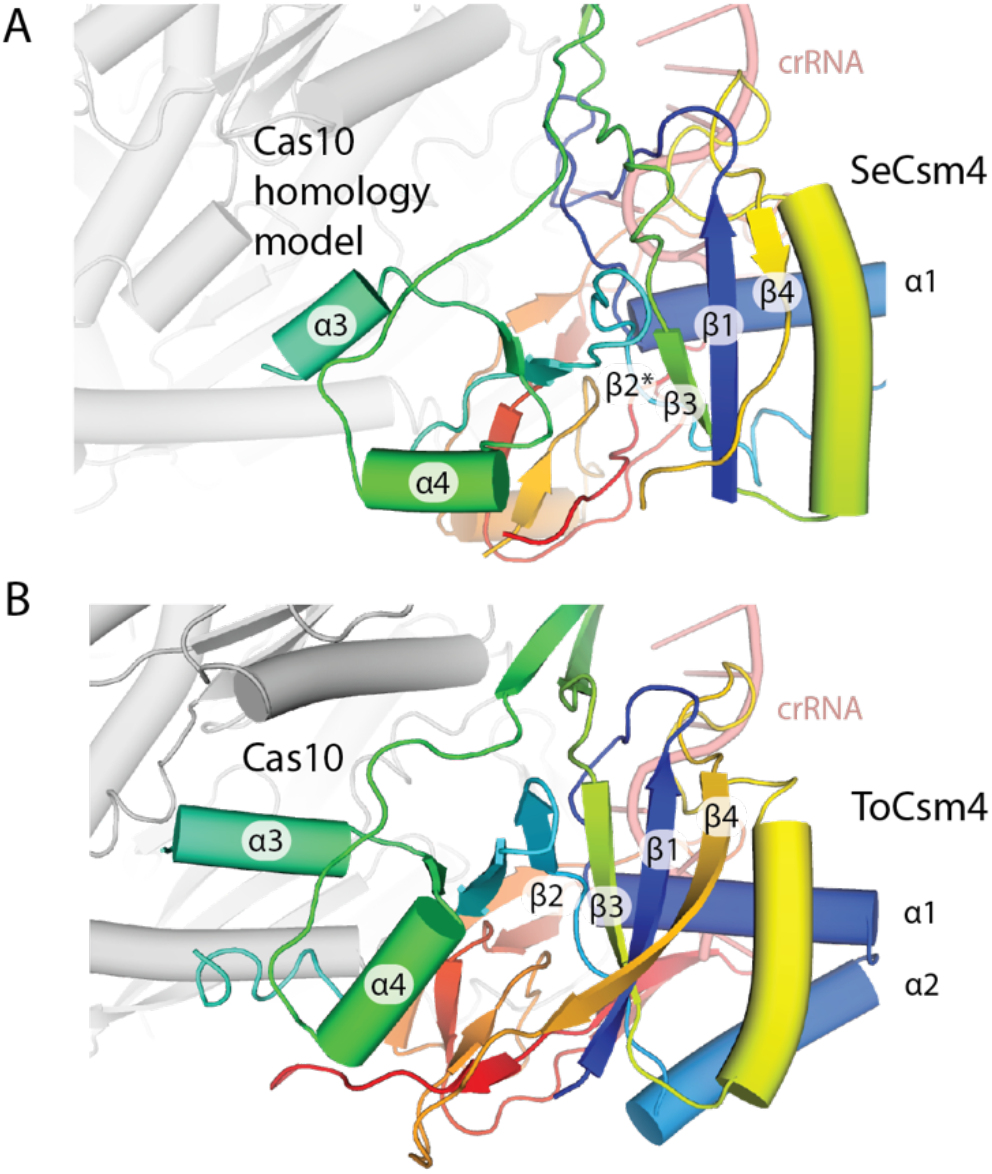
Csm4 possesses similar secondary structure elements in *T. onnurineus* and *S. epidermidis*. (*A*) The RNA recognition motif core of the Cas5 family member, SeCsm4, is elaborated to contact the 5’ tag of crRNA and Cas10. A homology model of SeCas10 is docked into the density present in *S. epidermidis* cryo-EM maps. (*B*) The secondary structure elements of ToCsm4 are similar to SeCsm4 but α3 has more extensive interactions with Cas10.

**Figure 7.**
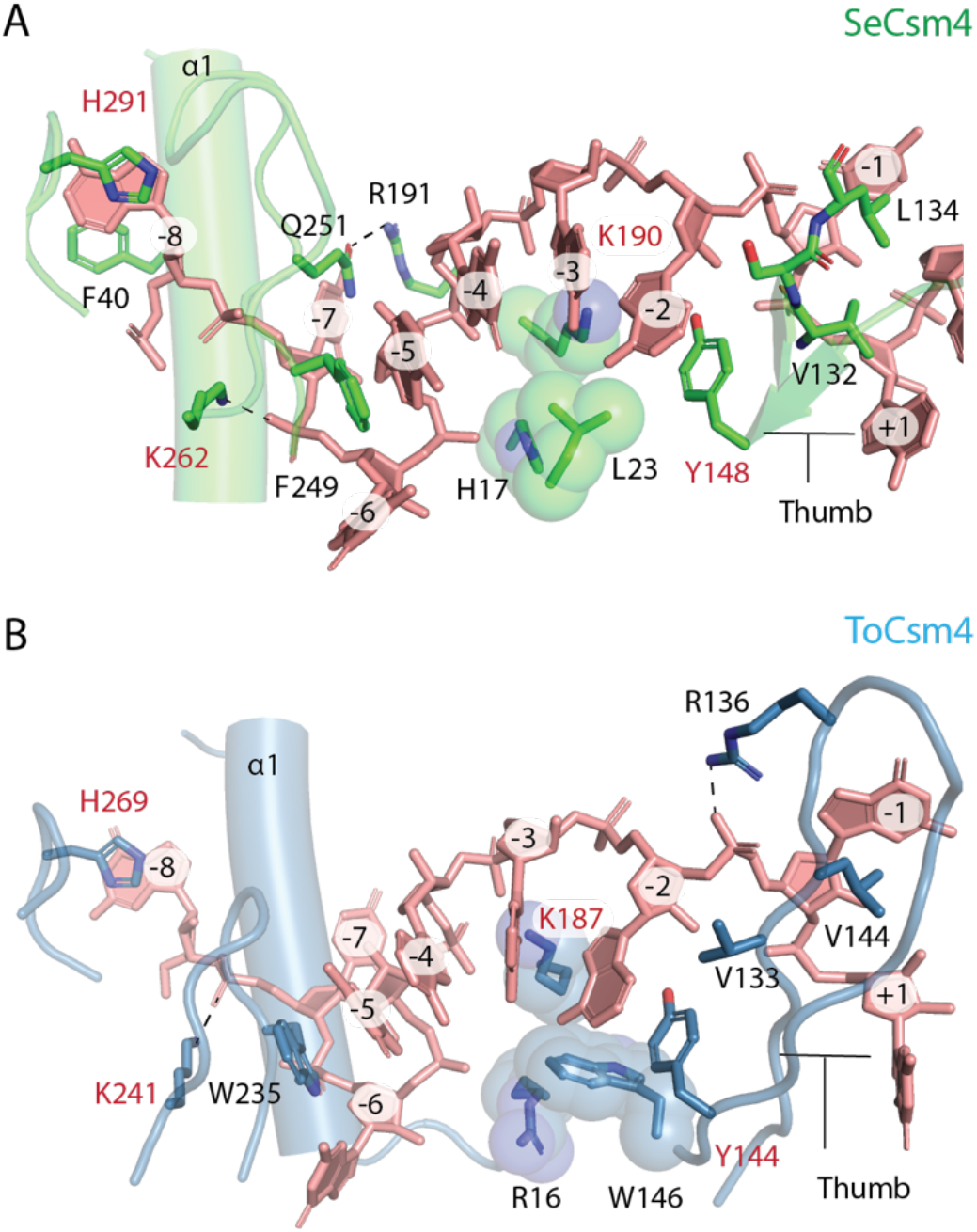
A detailed comparison of the interactions of SeCsm4 and ToCsm4 with crRNA. (*A*) SeCsm4 residues forming interactions with the crRNA 5’-tag that are likely critical for recognition shown as sticks. Three residues that form van der Waals interaction with crRNA and sterically position it are shown as spheres. Residues that are conserved between SeCsm4 and ToCsm4 are labeled in red. The thumb, β-hairpin-like element, that interrupts the co-axial nucleotide stack is labeled. The first α-helix of Csm4 is marked as α1. (*B*) ToCsm4 residues forming critical interactions with crRNA are shown as sticks or spheres. The β-hairpin-like element thumb is labeled as is α-helix 1, α1.

The pattern of interaction with crRNA is conserved between SeCsm4 and ToCsm4 and several critical residues are conserved, including the His that pi-stacks to the −8 nt, the Lys residue that sits beneath the −3 nt and the Tyr that pi-stacks with the −2 nt. An additional Lys residue, K241 in ToCsm4, is conserved between the archaeal and bacterial Csm4 however appears to interact with the −6 nt in SeCsm4 versus the −7 nt in ToCsm4 (Figs. 7A, 7B). A difference between the two Csm4 proteins is in the thumb region where ToCsm4 appears to use R136 to stabilize the flipped −1 nt. In contrast to Csm2 and Csm3 where comparison of archaeal and bacterial structures suggest likely mechanistic differences, the Csm4 structures suggest a conserved mode of interaction with crRNA.

Csm5 is a RAMP domain protein that binds the 3’ end of the crRNA-target duplex. The position of Csm5 in the complex suggests it blocks extension of the Csm3 and Csm2 oligomers and promotes nucleolytic maturation of the 3’ end of crRNA by cellular nucleases - two processes that are likely interrelated [50]. The RAMP domain of Csm5 blocks the extension of the Csm3 oligomer and the primarily α-helical capping domain blocks extension of the Csm2 oligomer (Figs. 1D, 1E and 8A). Five regions of Csm5 interact with RNA including the β1-loop, the capping domain, the thumb, α-helix 1 and the β3’-loop (Fig. 8A). The β1-loop possesses two conserved Gly residues allowing a sharp kink in the peptide backbone that hydrogren bonds to crRNA (Fig. S9A). The thumb interacts with the major groove of the RNA duplex but doesn’t intercalate the duplex as the Csm3 thumb does (Fig. 8B). The β3’-loop and α1 appear to work in unison gripping the crRNA in the vicinity of nucleotides +17 to +21 in the 276 kDa complex (Fig. 8C). Interestingly, two SeCsm5 acidic residues, D162 and E191, that were previously shown to be required for crRNA maturation are observed participating in electrostatic interactions (Fig. 8D) [50]. E191, which sits at the base of the thumb interacts with R121 that is positioned slightly C-terminal to α-helix 1. A potential explanation for the role of E191 in crRNA maturation is that the E191-R121 interaction allows cross-talk between the two crRNA binding regions, perhaps through an allosteric mechanism that enhances Csm5’s affinity for crRNA. The interaction between Csm5 residue D162 and the Csm3 residue R141 might contribute to crRNA maturation by influencing how Csm5 is positioned relative to its neighbors within the Cas10-Csm complex. Such positional and allosteric changes in Csm5 may impact the recruitment of PNPase and RNaseR, cellular nucleases which are now known to bind Csm5 and catalyze crRNA maturation [51, 52].

**Figure 8.**
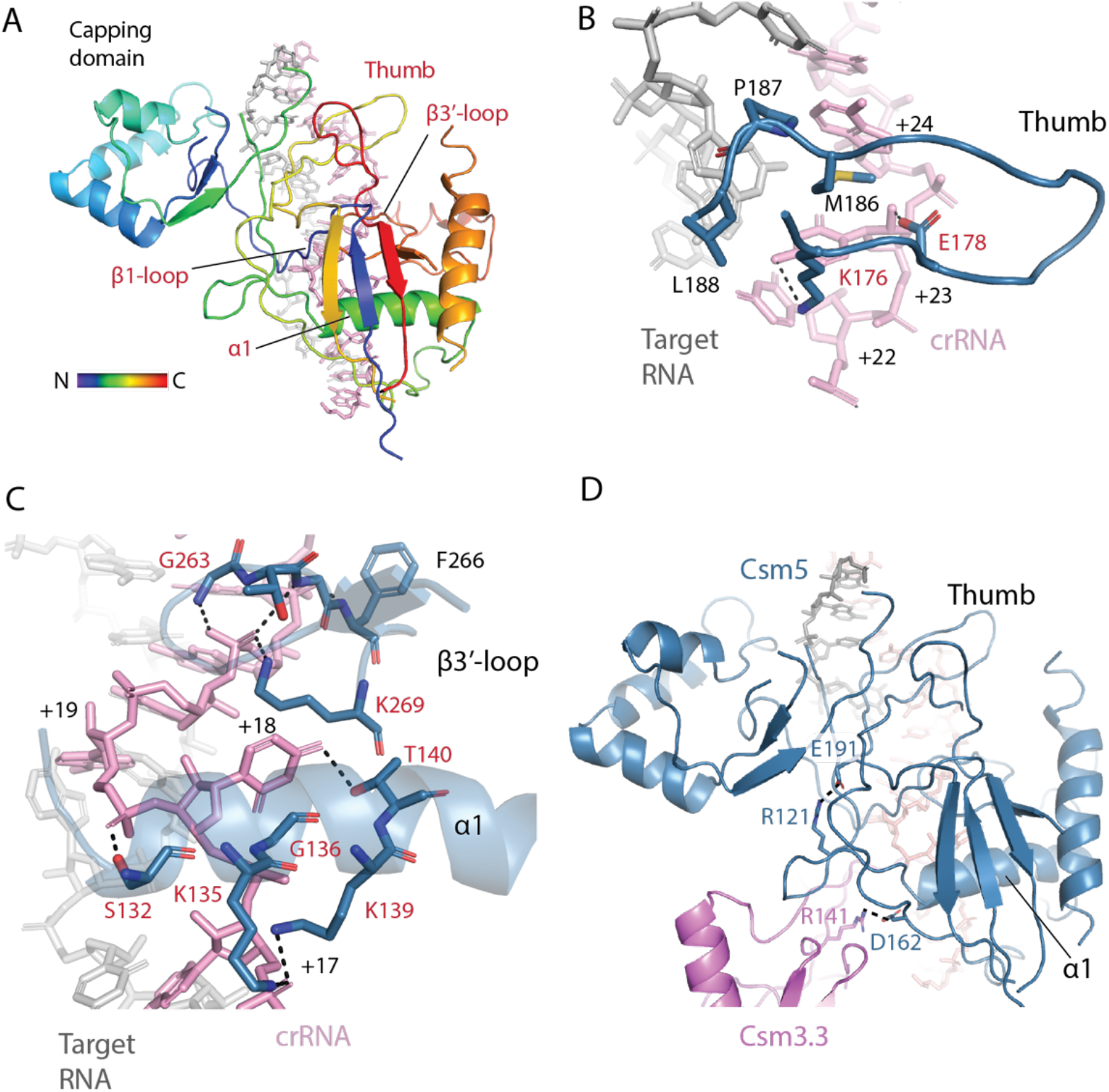
Most RNA-interacting regions of Csm5 are conserved and an electrostatic interaction may allow cross-talk between two regions. (*A*) SeCsm5 bound to the crRNA-target duplex is shown emphasizing regions that contact the duplex. Four of the five regions are conserved (red) across diverse Csm5 homologs. The secondary structure elements of Csm5 (β1, α1 and β3’) are named according to their position in the core, RRM domain - capping domain elements were bypassed. (*B*) The SeCsm5 Thumb region interacts with the major groove of the duplex by hydrogen-bonding and van der Waals interactions. Residues conserved across SeCsm5 homologs are colored red. (*C*) Helix-α1 and the β3’-loop interact with the duplex surrounding the flipped +18 nucleotide. (*D*) Electrostatic interactions between R121 and E191 may conformationally couple the helix-α1 and thumb components of SeCsm5. The interaction of SeCsm5 D162 with Csm3 R141 has been shown to be critical for crRNA maturation.

Structural superpositions and multiple sequence alignments indicate that four of the five Csm5 regions that interact with RNA are conserved from bacteria to archaea (Figs. 8, S9). The capping domain, however, that exclusively interacts with target RNA, diverges in structure and sequence even among bacteria (Fig. 9). SeCsm5 uses the basic residues K25 and K26 on the N-terminal end of the capping domain to form electrostatic interactions to target RNA (Figs. 9A, 9B). These residues show modest conservation among bacterial Csm5 proteins but the additional SeCsm5 contacts to target RNA, such as R71, E72 or N114, display even less conservation (Fig. 9A). Structural comparisons of SeCsm5 to StCsm5 confirm divergent contacts to target RNA are occuring and even show that StCsm5 possesses different secondary structure in the capping domain compared to SeCsm5 (See the arrow in Fig. 9C). Inspections of the ToCsm5 capping domain, suggests minimal contacts to target RNA in the archaeal Csm5 (Fig. 9D). In sum, while the RAMP domain of Csm5 possesses a pattern of interactions with crRNA consistent with that of other RAMP domain proteins (Csm3 and Csm4), the capping domain, which contacts target RNA, possesses idiosyncratic structures which suggest the possibility for functional differences among Csm5 proteins. Two potential areas for functional divergence could be in target RNA sensing or in the recruitment of cellular nucleases for crRNA 3’ end maturation [51, 52].

**Figure 9.**
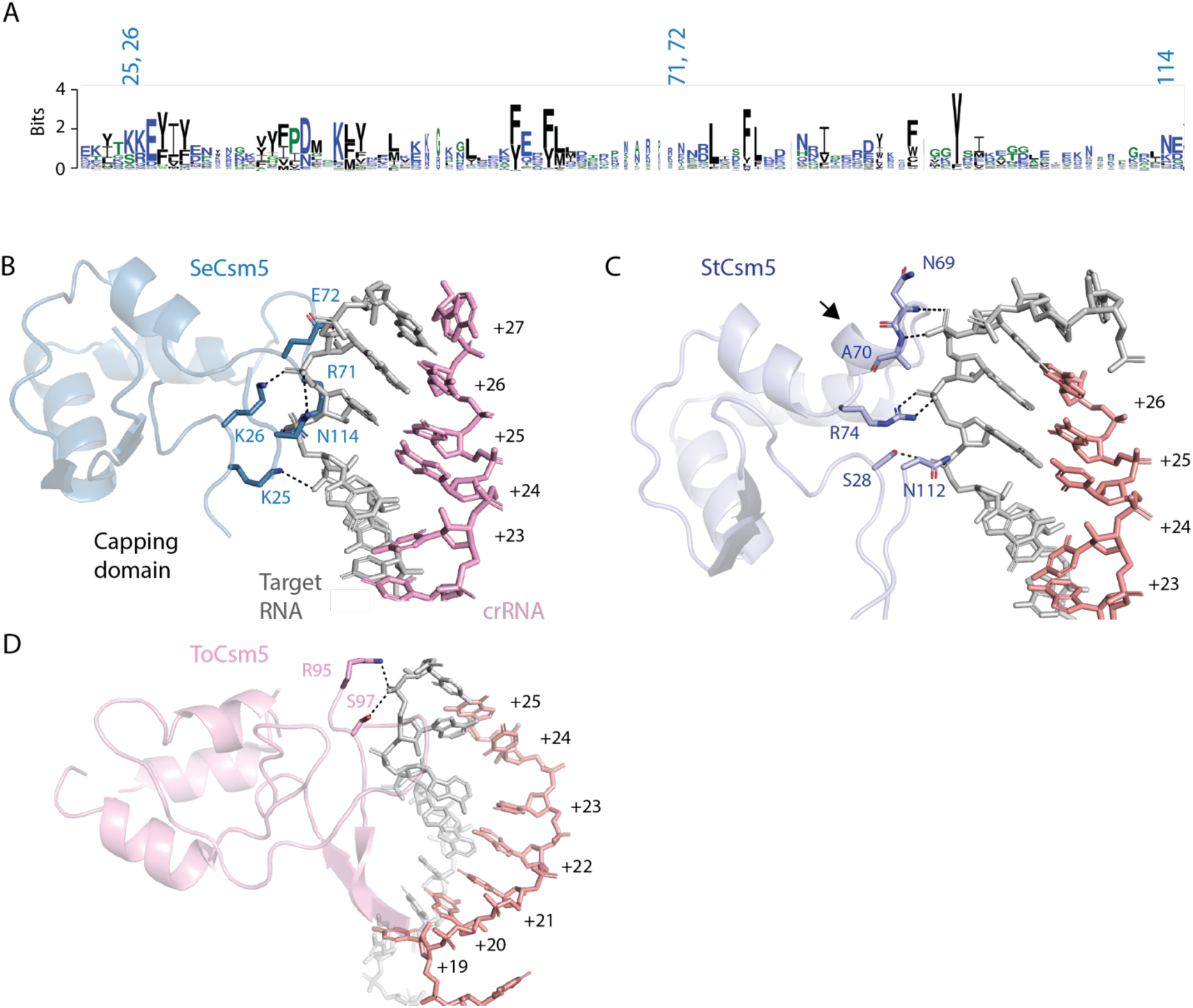
The capping domain of Csm5 diverges among Csm5 proteins. (*A*) A sequence logo of the capping domain with the positions of SeCsm5 residues that interact with target RNA numbered. The logo indicates there is limited sequence conservation within the capping domain. (*B*) Electrostatic and hydrogen bonding interactions between SeCsm5 and target RNA are indicated by dashed lines. (*C*) The interactions of the StCsm5 capping domain with target RNA are indicated. StCsm5 contains a helical segment, indicated by an arrow, absent in the other Csm5 structures. (*D*) Only two residues in the capping domain of the ToCsm5 structure reported in PDB code 6mus contact target.

## Discussion

We report cryo-EM data demonstrating that SeCas10-Csm forms two oligomerization states, a 276 kDa complex with the stoichiometry, Cas10_1_ Csm2_2_ Csm3_3_ Csm4_1_ Csm5_1_, and a 318 kDa complex that is formed by addition of an extra copy of Csm2 and Csm3. These two stoichiometries were recently observed in the cryo-EM reconstruction of *L. lactis* Cas10-Csm [21]. Recent cryo-EM reconstructions of SeCas10-Csm by Smith and co-workers (published while this manuscript was in preparation) reported these two stoichiometries as well and observed a Cas10_1_ Csm4_1_ Csm3_5_ stoichiometry; five copies of Csm3 were observed and Csm2 and Csm5 were not present [24]. Biochemical assays of SeCas10-Csm by Smith and co-workers demonstrated the presence of Csm2 and Csm5 in the as-isolated complex and the complexes were purified from *S. epidermidis* cells as ours also were. It is likely that in the absence of target RNA and under the conditions Smith and co-workers used for EM grid preparation, Csm2 and Csm5 weakly associate with the complex [24]. A weak association of Csm2 with apo *L. lactis* Cas10-Csm has been reported and this phenomenon is likely linked to conformational dynamics critical for regulating interference that will be discussed below [21]. The Csm3_n_ Csm2_n-1_ stoichiometry, that we report, has been observed for Cas10-Csm complexes from other species as well [20–22]. The existence of two stoichiometries of SeCas10-Csm differing by the presence of one Csm2-Csm3 unit was anticipated due to two previous observations: in the presence of a crRNA, in vitro, Csm3 assembles into oligomers with protomer number depending on crRNA length and as-isolated SeCas10-Csm contains primarily two lengths of crRNA, a 37-mer and 43-mer (Fig. 1C) [27].

The identification of the two SeCas10-Csm states, 276 kDa and 318 kDa, is important because it has been recently argued that the co-existence of multiple oligomerization states in Type III CRISPR complexes imparts a unique interference behavior. Several researchers have shown that cOA synthesis is sensitive to mismatches in base-paring between target and crRNA proximal to Cas10, the region denoted as positions +1 to +7 or the Cas10-activating region (Fig. 1B) [12–14, 29]. Steens and colleagues argue that the region of the target-crRNA duplex distal to Cas10, composed of positions +19 to +37, also performs a check on base-pairing complementarity by controlling initiation of duplex formation [13]. Mismatches in the vicinity of the 3’ end crRNA disfavor target binding and thus interference. However, this effect is obscured when assayed in a population of mixed oligomerization states since the complexes contain different crRNA 3’ ends [13]. The interaction of these phenomenon suggests that distinct oligomerizatoin states of Type III CRISPR complexes are sensitive to mismatches at both the 5’ and 3’ end of crRNA but the effect of mismatches at the 3’ end are obscured when pooled oligomerization states are assayed. The data indicating base-pairing at the crRNA 3’ end critically controls interference was obtained in a Type III-B complex which contains the Cmr1 protein at this region, a protein absent from Type III-A complexes. Therefore an important future direction is to determine the precise mechanistic effects of mismatches throughout the crRNA-target duplex in Type III-A CRISPR and determine how Cas10-Csm oligomerization state affects sensitivity to mismatches. A complete understanding of how mismatches effect cOA synthesis in Type III CRISPR is required to understand the contributions of these complexes to bacterial physiology and to efficiently develop molecular diagnostics based on them.

The importance of Csm2 in Cas10-Csm mediated interference has been questioned because in vitro experiments showed that the critical biochemical activities remain in a *S. thermophilus* complex lacking Csm2 [53]. However, later experiments with *S. thermophilus* Cas10-Csm complex containing a Csm2 site-directed mutant showed a complete loss of target RNA cutting and experiments with *L. lactis* complexes showed severe defects in DNase activity, cleavage of target RNA and cOA synthesis in a complex lacking Csm2 [20, 21]. Importantly, in vivo experiments in *S. epidermidis* and *L. lactis* revealed that interference falls to background levels in the absence of Csm2 [21, 54]. A caveat of these studies is that deletion of Csm2 caused a depletion of Csm5 in the complex and the interference defects could be the combined effect of depletion of both proteins.

Csm2 contacts to cognate target RNA were well-resolved in our 3.1 Å SeCas10-Csm reconstruction. These contacts could not be resolved in previous reconstructions of SeCas10-Csm due to the lower resolution of those structures [24]. Interactions of Csm2 residues along α-helix 2 with target RNA are a common feature among structures from multiple organisms (Fig. 3C) [20, 21, 23]. In the *S. thermophilus* complex Csm2 residue R41, which is structurally equivalent to SeCsm2 residue R49, has been shown to be required for target RNA cleavage [20]. A similar finding was made in *L. lactis* where Csm2 residue R48 is required for target RNA cleavage and the authors argue R48 acts as a general base on the 2’ hydroxyl of the labile nucleotide in an RNAse A like mechanism (Fig. 5B) [21]. It is then unsurprising that this Arg is the most conserved residue in α-helix 2 (Fig. 3B). While multiple studies observe interactions of Csm2 with target RNA similar to ours, the *T. onnurineus* Cas10-Csm structure diverges substantially (Fig. 3D). ToCsm2 residue R96, which is equivalent to SeCsm2 R49 in multiple sequence alignments, does not contact the 2’ hydroxyl of the labile, target nucleotide raising the question of why this residue is conserved in ToCsm2. One possibility is that the thermophilic ToCas10-Csm complex has not adopted its final, target bound conformation in the available cryo-EM reconstruction, a conformation that would involve a reorganization of ToCsm2 so that R96 contacts the labile target RNA residue [22].

Cas7 family members play a critical role in the function of Type I and Type III CRISPR systems by forming an oligomer that binds crRNA and presents it to solvent for base-pairing [48]. In Type III CRISPR systems, the Cas7 family member, Csm3, plays an additional role providing an Asp residue to the active site that cleaves target RNA [8, 55]. The RNase loop containing the Asp residue binds in the major groove of the crRNA-target duplex but sits near the 2’OH of the labile nucleotide because this nucleotide is flipped out of the co-axial stack (Figs 5A, 5B). Structures of Cas10-Csm from *T. onnurineus* and *L. lactis* were determined using Asp to Ala mutants of the critical residue to prevent target RNA cleavage during structure determination [21, 22]. We used the alternate approach of incubating the complex with EDTA to chelate Mg^2+^ allowing determination of a structure with wild type sequence. You and co-workers used a similar approach determining the structure of *S. thermophilus* Cas10-Csm with wild type sequence [20]. In both the structures with wild type sequence the Asp residue is pointed towards the 5’O of the reaction product at a distance of 4-5 Å (Fig. 5B). The recent structure of SeCas10-Csm bound to cognate target (non-self) RNA by Smith and co-workers observed a distance of 10.2 Å between Csm3 D32 and the labile phosphate [24]. This large distance is explained by the fact that the investigators captured a product complex with cleaved target RNA. Sridhara and co-workers have proposed an RNase A-like mechanism for target RNA cleavage and the position of the critical Asp is consistent with its role as the general acid in this mechanism but does not explain the importance of Mg^2+^ for target cleavage [21]. Future studies are needed to clarify the mechanism of target RNA cleavage but structural and biochemical data are consistent in implicating SeCsm2 R49 and SeCsm3 D32 as the critical residues for this activity [8, 20, 21, 55].

Our analysis of Csm3 interactions with the minor groove of the crRNA-target duplex revealed a major difference between archaeal and bacterial Csm3 (Fig. 5C). Archaeal Csm3 engages in interactions with the target RNA in the minor groove but the bacterial protein does not. Additionally, a Zn^2+^ ion binds adjacent to α2 and α2’ in the archaeal Csm3 but not in the bacterial protein. The functional siginficance of Csm3 interactions with the minor groove are not known but it seems reasonable to think they play a role in sensing bound target RNA, including a role in the sensitivity of Cas10-Csm to mismatches in base pairing. If this is the case, the differences in archaeal and bacterial Csm3 may mediate differences in mismatch sensitivity and there could be a role for the Zn^2+^ ion in target RNA detection in archaeal Cas10-Csm.

Functional studies have implicated a role for Csm5 in the maturation of the 3’ end of crRNA [50], and it is now understood that Csm5 recruits two cellular nucleases, PNPase and RNaseR, which catalyze crRNA maturation [51, 52]. Site-directed mutants of two acidic residues in SeCsm5 cause defects in crRNA maturation but the mechanism for this effect was unclear. Our structural results suggest an explanation. The R121-E191 interaction may allow cross-talk between the two cRNA binding regions of Csm5, the thumb region and the α-helix 1 region (Fig. 8D). For example, binding of the α-helix 1 region to crRNA could influence nearby R121 to promote the formation of the electrostatic interation with E191 enhancing the affinity of the thumb region for crRNA. While E191 has only been functionally investigated in *S. epidermidis*, this residue is conserved in Csm5 homologs and in the *S. thermophilus* structure reported by PDB code 6ifu the residue and its electrostatic interaction are conserved (Fig. S9D). Therefore, the R121-E191 interaction may modulate Csm5 conformational changes and thus the recruitment of crRNA maturation nucleases in organisms harboring Type III systems. Our structure revealed that SeCsm5 residue D162 forms an electrostatic interaction with a Csm3 residue (Fig. 8D). We hypothesize that the crRNA maturation defect arising from D162A is caused by a mis-positioning of Csm5 relative to Csm3 in this mutant. Further studies are needed to investigate this but it is notable that the D162-Csm3 R141 electrostatic interaction is also conserved in *S. thermophilus* (Fig. S9D).

Csm5 has been found to influence the affinity of target RNA binding to *S. thermophilus* Cas10-Csm and in a Type III-B system it was found that mismatches between the 3’ end of cRNA and target decreased affinity for target RNA leading to interference defects [13, 53]. Both results suggest an important role for Csm5, or its Type III-B homolog Cmr6, in Type III CRISPR function. Importantly, we note in our structural analyses that the only region of Csm5 in Type III-A complexes that contacts target RNA is the capping domain, a region that diverges in structure and sequence among Csm5 homologs (Fig. 9). Additionally, substantial conformational change in the Csm2 subunits have been observed upon target RNA binding to StCas10-Csm [20]. Since the capping domain of Csm5 contacts both target RNA and Csm2, we speculate that there may be an underlying connection between the phenomenon mentioned above: that interaction of the capping domain of Csm5 with target RNA promotes conformational changes in Csm2 that enhance its affinity for target RNA, potentially a critical early step in activating interference. The fact that the Csm5 capping domain diverges in structure and sequence appears to conflict with the argument that it plays a crucial role in interference, however, it may be that differences in the capping domain are necessary for Csm5 to interface with non-Cas cellular nucleases sepcific to each organism.

Type III CRISPR systems are uniquely suited to serve as molecular diagnostics because these systems are adapted to perform specific detection of nucleic acids and amplify the detection event by a multiple-turnover enzymatic assay. This intrinsic potential of Type III CRISPR systems was realized in the last year with the publication of four examples of the deployment of a Type III complex to detect viral RNA [11–14]. We believe that the subtle variations in the structure of Type III-A complexes that we have described suggest there may be activity differences in these complexes, such as their intrinsic sensivity to mismatches in base-paring or affinity for target RNA, that would make one complex better suited to a role in molecular diagnostics than another. Additionally, we confirm that SeCas10-Csm forms two oligomeric states, a 276 kDa complex and a 318 kDa complex, and we believe this information will aid in investigating how mismatches at the 3’ end of crRNA-target duplex affect interference, a question whose answer is important in the construction of molecular diagnostics and other biotechnologies based on Type III CRISPR-Cas systems.

## Data availability

The atomic coordinates, PDB code 8DO6, have been deposited with the Protein Data Bank. Maps derived by single-particle reconstructions using cryo-EM have been deposited with the Electron Microscopy Data Bank, emd-27593 (276 kDa complex) and emd-27762 (318 kDa complex).

## Author contributions

M.P., M.N., L.C-Z., S.A.K., A.J.S. and J.A.D investigation; A.J.S, A.H-A., S.M.S and J.A.D supervision; M.P., A.J.S, A.H-A., S.M.S and J.A.D writing-reviewing and editing. A.J.S, A.H-A., S.M.S and J.A.D funding acquisition.

## Funding and additional information

This work was supported by NIGMS, National Institutes of Health Grant R35GM142966 (to J. A. D.). NIH grants GM103310 and GM139171, and the Simons foundation grant SF349247 (to NYSBC). Single particle data collection was done at Simons Electron Microscopy Center at New York Structural Biology Center. Thanks to Misha Kopylov for preparing the microscope for data collection. SEC-SAXS work was conducted at the Advanced Light Source (ALS), a national user facility operated by Lawrence Berkeley National Laboratory on behalf of the Department of Energy, Office of Basic Energy Sciences, through the Integrated Diffraction Analysis Technologies (IDAT) program, supported by DOE Office of Biological and Environmental Research. Additional support comes from the National Institute of Health project ALS-ENABLE (P30 GM124169) and a High-End Instrumentation Grant S10OD018483.

