## Supporting_Information for "The structure of a Type III-A CRISPR-Cas effector complex reveals conserved and idiosyncratic contacts to target RNA and crRNA among Type III-A systems"

**Table S1. Oligonucleotides used in the study.**

| Name | Sequence (5'-3') | Description |
|---|---|---|
| F063 | TTGCTGCTTAATATATTGCATCATCAAAGATAAACC | Gibson assembly, p*crispr-spc1* |
| A010 | CTTTGTACTGATGATTTATATACTTCGGCATACG | Gibson assembly, p*crispr-spc1* |
| F062 | TTATCTTTGATGATGCAATATATTAAGCAGCAAGAG | Gibson assembly, p*crispr-spc1* |
| L162 | CGAAGTATATAAATCATCAGTACAAAGTAAATCTAACAACACTCTAAAAAATTGTAGATTTTGAATAAAATACG | Gibson assembly, p*crispr-spc1* |
| A200 | TTGTCAAAAAAAGTGACATATCATATAATCTTGTAC | Sequencing confirmation of p*crispr-spc1* |
| F111 | GCGTCCACGTTTAAATTGTTTGC | Sequencing confirmation of p*crispr-spc1* |
| ssRNA-01 | CUUUGUACUGAUGAUUUAUAUACUUCGGCAUACGUUCUCUAAA | Analog of Nickase mRNA, cognate target for Spc1 crRNA |

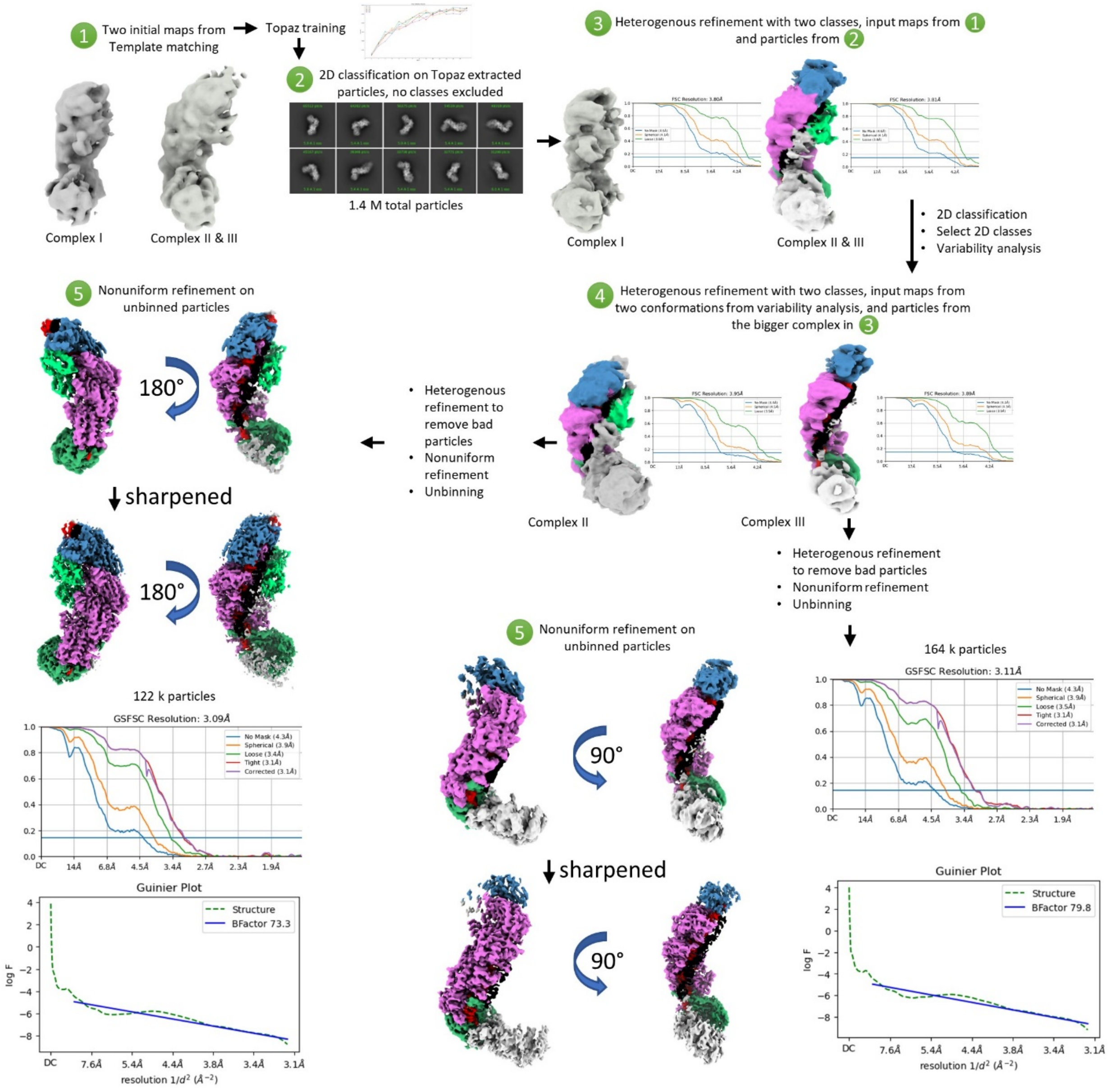


**Figure S1. Processing workflow in cryoSPARC V2.** At step 1, two initial maps with different stoichiometries were obtained using template matching. From particles making those maps a set of 5000 particles were used for Topaz training at step 2. Complexes I, II, and III denote the smallest to largest stoichiometries. At step 3, heterogenous refinement was done on the full set of 1.4 million particles. Cas10 is grey, Csm2 is light green, Csm3 is magenta, Csm4 is dark green and Csm5 is blue. At step 4, the map denoted as complex II and III was further classified with two input maps from variability analysis. Each of the complexes II and III were refined to high resolution with unbinned images at step 5. FSC curves and B-factor plots are included. Due to flexibility of the Cas10, this density is averaged out in the high-resolution refinement of complex II and to a lesser extent in complex III. Csm2 densities are also averaged out at high resolution in complex III because they were under-populated. Complex II was used for the molecular models presented in this manuscript.

**Table S2. Components of the atomic model of SeCas10-Csm (276 kDa complex) bound to target RNA**

| Name | Uniprot code or sequence (5’-3’) | Description |
|---|---|---|
| Csm2 | Q5HK90 | Csm2 protein sequence from *S. epidermidis* RP62A |
| Csm3 | Q5HK91 | Csm3 protein sequence from *S. epidermidis* RP62A |
| Csm4 | Q5HK92 | Csm2 protein sequence from *S. epidermidis* RP62A |
| Csm5 | Q5HK93 | Csm5 protein sequence from *S. epidermidis* RP62A |
| crRNA | ACGAGAACACGUAUGCCGAAGUAUAUAAAU CAUCAGU | crRNA derived from *spc1* of *S. epidermidis* RP62A |
| ssRNA-01 | CUUUGUACUGAUGAUUUAUAUACUUCGGC AUACGUUCUCUAAA | Analog of *nickase* transcript, synthetic oligonucleotide |

**Table S3. SAXS rigid body components**

| Rigid body unit | 6ifu components (chain ID) | rigid body constraints |
|---|---|---|
| 1 | Csm1 (A) K2 – K654, Csm4 (G), Csm3 (F), crRNA (I) A1 – U13, CTR2 RNA (J) A30 – C34 | 4.0 A between unit 1 (Csm1 K654) and unit 2 (Csm1 F655), 6.5 A between unit 1 crRNA U13 and unit 2 crRNA C14, 6.5 A between target RNA A30 and unit 2 target RNA G29 |
| 2 | Csm1 (A) F655 – K757, Csm3 (E), crRNA (I) C14 – U19, CTR2 RNA (J) A24 – G29 | 6.5 A between unit 2 crRNA U19 and unit 3 crRNA C20, 6.5 A between unit 2 target RNA A24 and unit 3 target RNA G23 |
| 3 | Csm2 (C), Csm3 (D), crRNA (I) C20 – U25, CTR2 RNA (J) A18 – G23 | 6.5 A between unit 3 crRNA U25 and unit 4 crRNA C26, 6.5 A between unit 3 target RNA A18 and unit 4 target RNA U17 |
| 4 | Csm2 (B), Csm5 (H), crRNA (I) C26 – A34, CTR2 RNA (J) A7 – U17 | |

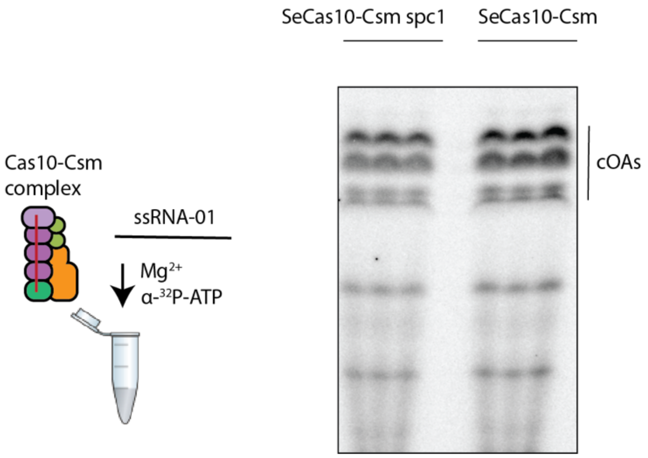


**Figure S2. Cyclic oligoadenylate synthesis by SeCas10-Csm encoded by p*crispr-spc1* or p*crispr*.** A 24% urea-PAGE gel was used to visualize production of $^{32}P$-containing cyclic oligoadenylates produced by the incubation of target RNA (ssRNA-01) and ATP with *S. epidermidis* Cas10-Csm expressed from the p*crispr spc1* (SeCas10-Csm spc1) plasmid which contains only one spacer gene or SeCas10-Csm expressed from the p*crispr* plasmid which contains all three spacer genes found in the *S. epidermidis* genomic, CRISPR locus . The products from three technical replicates are shown.

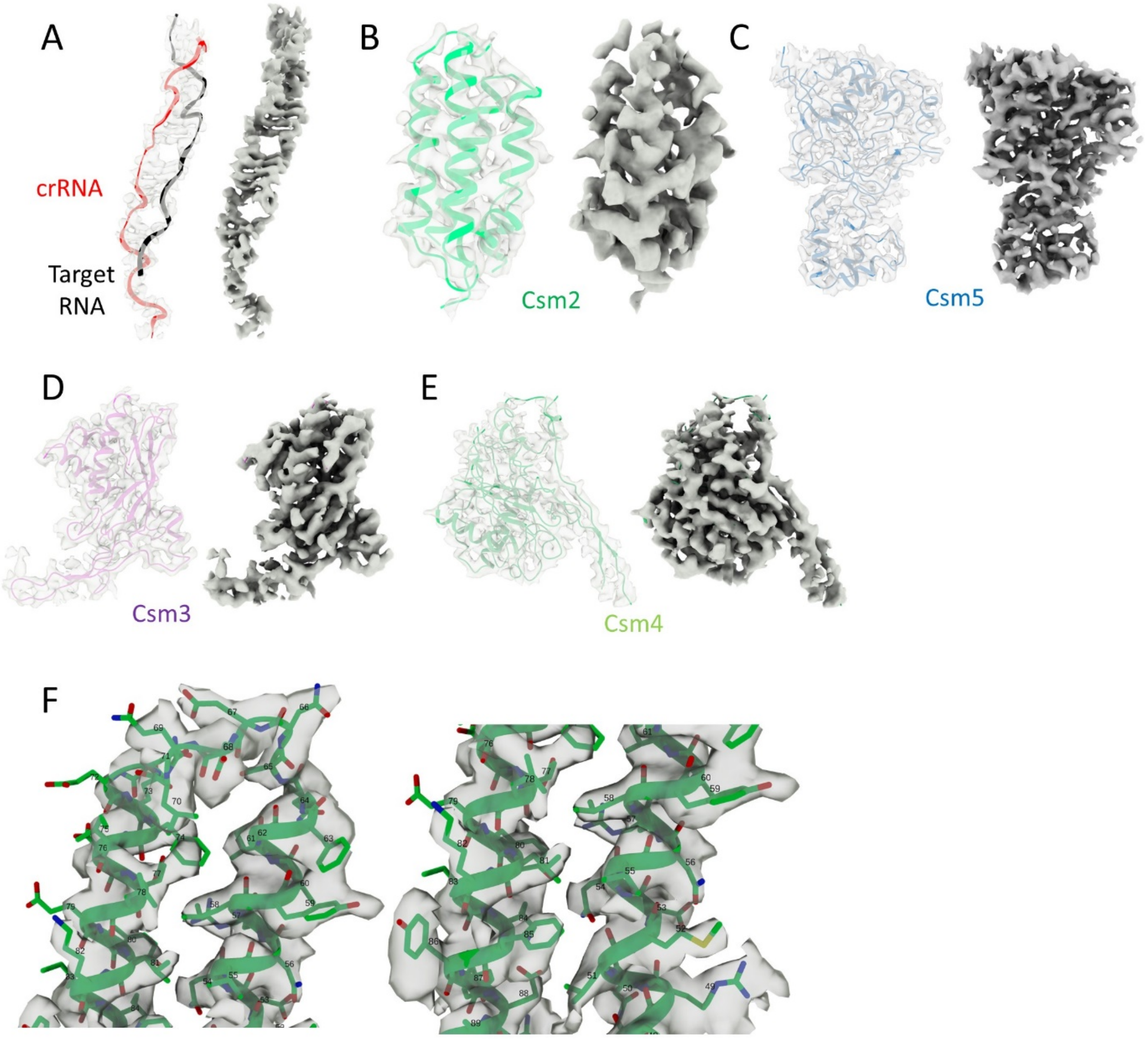


**Figure S3. Cryo-EM map quality.** Different subunits of the Cas10-Csm complex are shown in each panel. Each panel contains the density on the right and the model fit in the density on the left. Panel F shows the side chain densities for Csm2 along with the residue numbers.

**Table S4. Cryo-EM Statistics for Data Collection and Model Quality**

| Data collection and processing | Cas10-Csm bound to target RNA (8DO6, 276 kDa complex) |
|---|---|
| Magnification | 81,000 |
| Voltage (kV) | 300 kV |
| Electron exposure ($e^-$/ Å$^2$) | 44.7 |
| Defocus range (μm) | 1.0-1.5 |
| Pixel size (Å) | 0.846 |
| Initial particles (no.) | 1,400,000 |
| Final particles (no.) | 122,000 |
| Map resolution (Å) | 3.1 |
| FSC threshold | 0.143 |
| Map sharpening B factor (Å$^2$) | 73.3 |
| Refinement<br>Csm2-5, target and crRNA | |
| Model resolution (Å) | 3.1 |
| FSC threshold | 0.143 |
| Model composition | |
| Nonhydrogen atoms | 12980 |
| Protein residues | 1450 |
| RNA residues | 61 |
| Bonds (RMSD) | |
| Bond lengths (Å) | 0.005 |
| Bond angles (°) | 0.679 |
| Validation | |
| Molprobity score | 2.16 |
| Clashscore | 16.50 |
| Ramachandran plot (%) | |
| Outliers | 0.42 |
| Allowed | 6.33 |
| Favored | 93.25 |
| B factors, mean (Å$^2$) | |
| Protein | 60.60 |
| RNA | 54.33 |

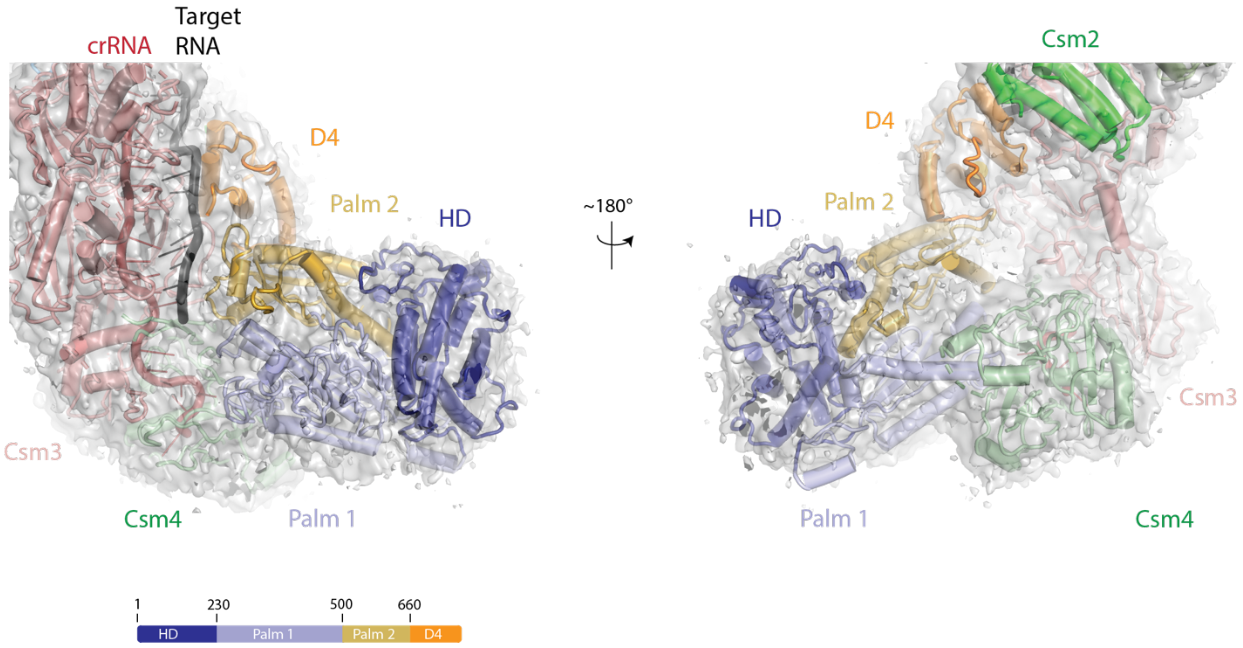


**Fig. S4. Density for each domain of Cas10 exists in the map of the 276 kDa Cas10-Csm complex.** A homology model of *S. epidermidis* Cas10 was docked into the density given by the map of the 276 kDa Cas10-Csm complex. The map is shown at σ=2.0. Cas10 is color coded by domain: HD, HD nuclease domain, Palm1, Palm1 polymerase domain 1, Palm 2, Palm polymerase domain 2, D4, domain 4.

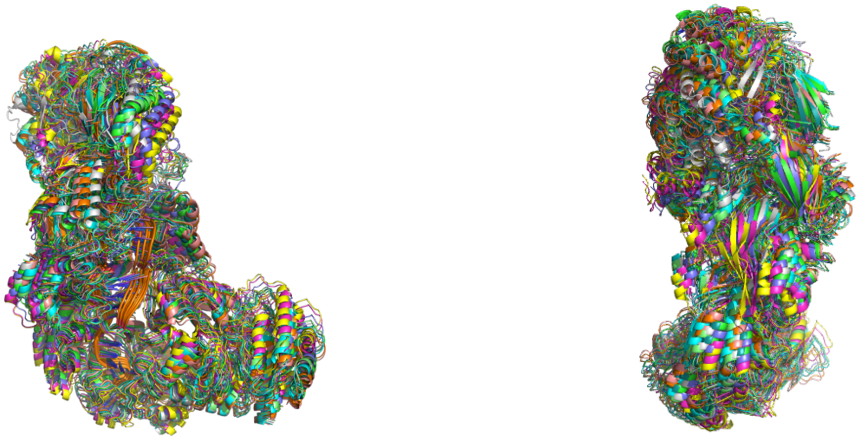

**Figure S5. Independently calculated rigid body models.** Ten rigid body models calculated in SASREF are shown overlaid with each other; the two views are related by 90 degrees. The overall arrangement and shape of the models are highly consistent.

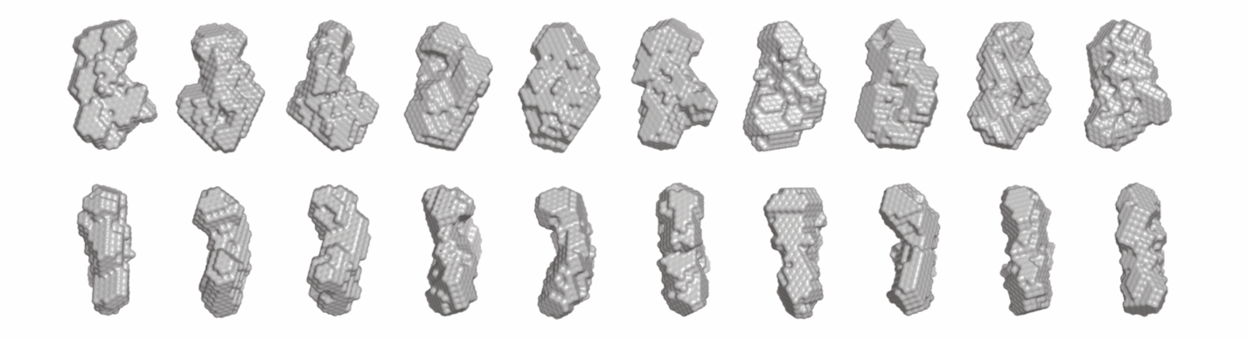

**Figure S6. Independently calculated *ab initio* models.** *Ab initio* models were calculated using DAMMIN, with a $D_{max}$ of 160. The upper and lower panels show the same models rotated by 90 degrees.

**Table S5. Small Angle X-ray Scattering Model Calculation Statistics**

| **Model type** | **No. of models** | **Average Chi$^2$** | **Best Chi$^2$** | **Average NSD or RMSD (in Å)** |
|---|---|---|---|---|
| rigid body | 10 | 1.724 ± 0.02 | 1.690 | 4.6 ± 2.5 |
| ***ab initio*** | 10 | 1.479 ± 0.001 | 1.477 | 0.603 ± 0.02 |

Sets of *ab initio* models were calculated from the experimental SAXS scattering profile using DAMMIN. Sets of rigid-body molecular models were calculated from the experimental SAXS scattering profile using SASREF (see Methods for details). Agreement of these models with the SAXS data are indicated by the Chi$^2$ values (calculated by DAMMIN for *ab initio* models and by SASREF for rigid body models); agreement of these models with each other are indicated with RMSD values (calculated in PyMOL) for molecular models and normalized spatial discrepancy (NSD) values (calculated in SUPCOMB) for *ab initio* models. The average RMSD for molecular models relatively high due to small positional shifts between domains, but the overall shape and architecture remains very similar (see Figure S5).

**Table S6. Agreement between SAXS *ab initio* models and molecular models**

| Molecular model type | No. of models | Average NSD | Best NSD |
|---|---|---|---|
| SAXS rigid body models | 10 | 1.8 ± 0.1 | 1.5909 |
| 276 kDa EM | 1 | 1.9 ± 0.1 | 1.6798 |
| 318 kDa EM | 1 | 2.1 ± 0.1 | 1.9360 |
| 318 kDa EM-no Csm2 | 1 | 2.5 ± 0.2 | 2.3224 |
| PDB ID 6IFU | 1 | 2.0 ± 0.1 | 1.7445 |

The indicated molecular models were superposed with the set of ten SAXS *ab initio* models using SUPCOMB (Kozin 2001), and the normalized spatial discrepancies (NSDs) from these superpositions were averaged. For the set of SAXS rigid body models, a comprehensive set of pairwise superpositions was performed.

**Table S7. CRYSOL fitting statistics**

| Models | $Chi^2$ |
|---|---|
| 6ifu | 1.568 |
| 276 kDa EM | 1.591 |
| 318 kDa EM | 2.657 |
| 318 kDa-noCsm2 EM | 3.017 |

The indicated models were used to generate theoretical scattering curves, which were then fit to the experimental SEC-SAXS scattering data using CRYSOL.

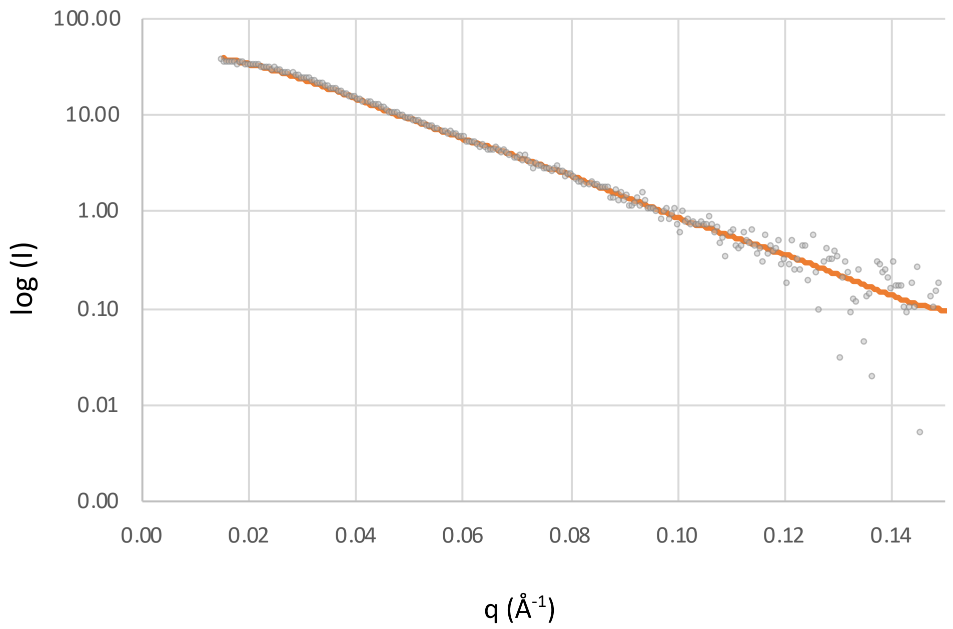


**Figure S7. Deconvolution.** EM models for the 276 kDa complex, the 318 kDa complex, and the 318 kDa complex without Csm2 were used to generate theoretical scattering curves, which were then used in OLIGOMER to deconvolute our experimental SAXS scattering curve. The resulting deconvolution strongly indicated that the 276 kDa complex alone was the best model for the experimental data. The composite fit is shown above as a line with the experimental data as open circles. The $Chi^s$ for this fit was 1.75.

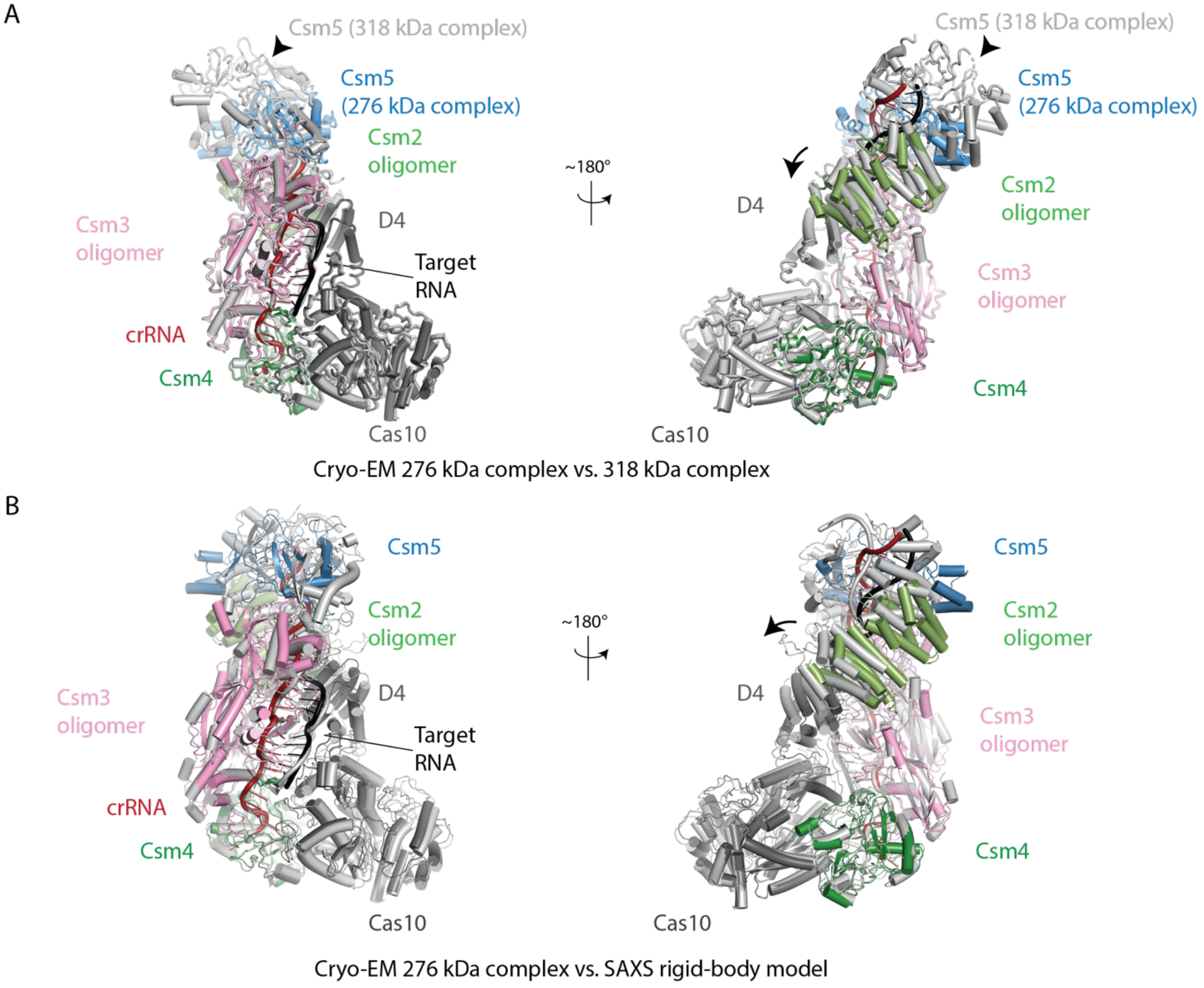


**Figure S8. Superposition of SeCas10-Csm models derived from cryo-EM and SAXS.** (*A*) Superposition of the 276 kDa SeCas10-Csm complex (multi-colored, PDB code 8DO6) and the 318 kDa SeCas10-Csm complex (8DO6 chains docked to density, grey) reveals they differ in the number of Csm2 and Csm3 subunits and possess a modest shift of the Csm2 oligomer towards Cas10 (arrow). (*B*) A superposition of the 276 kDa molecular model (multi-colored, PDB code 8DO6) with a rigid-body model derived by SAXS is shown. Again, a modest shift of the Csm2 oligomer towards Cas10 is observed (arrow). Superpositions were performed using Csm4. D4, refers to Cas10 domain 4, the C-terminal domain of the protein.

**Table S8. Sequence identities (% identity) among structurally characterized Cas10-Csm complexes.**

| | | | Ll | St | To |
|---|---|---|---|---|---|
| **Cas10** | Se | Q5HK89 | 48.1 | 36.1 | 23.9 |
| | Ll | L0CEJ3 | | 38.8 | 23.8 |
| | St | 6ig0_A | | | 23.2 |
| | To | B6YWB8 | | | |
| **Csm2** | Se | Q5HK90 | 33.3 | 28.9 | 14.5 |
| | Ll | L0CFW2 | | 34.6 | 16.5 |
| | St | 6ig0_C | | | 23.3 |
| | To | B6YWB9 | | | |
| **Csm3** | Se | Q5HK91 | 53.8 | 48.1 | 34.1 |
| | Ll | L0CEA3 | | 53.5 | 30.9 |
| | St | 6ig0_E | | | 32.8 |
| | To | B6YWC0 | | | |
| **Csm4** | Se | Q5HK92 | 46.1 | 31.7 | 22.4 |
| | Ll | L0CFH1 | | 35.4 | 21.5 |
| | St | 6ig0_G | | | 19.1 |
| | To | B6YWC1 | | | |
| **Csm5** | Se | Q5HK93 | 35.2 | 24.2 | 12.0 |
| | Ll | L0CG31 | | 25.0 | 15.5 |
| | St | 6ig0_H | | | 9.6 |
| | To | B6YWC5 | | | |

Se, *Staphylococcus epidermidis*. Ll, *Lactococcus lactis*. St, *Streptococcus thermophilus.* To, *Thermococcus onnurineus*. Uniprot codes are given for each protein except *S. thermophilus* strain ND03 sequences which are not present in Uniprot. Pairwise sequence alignments were made with Clustal Omega.

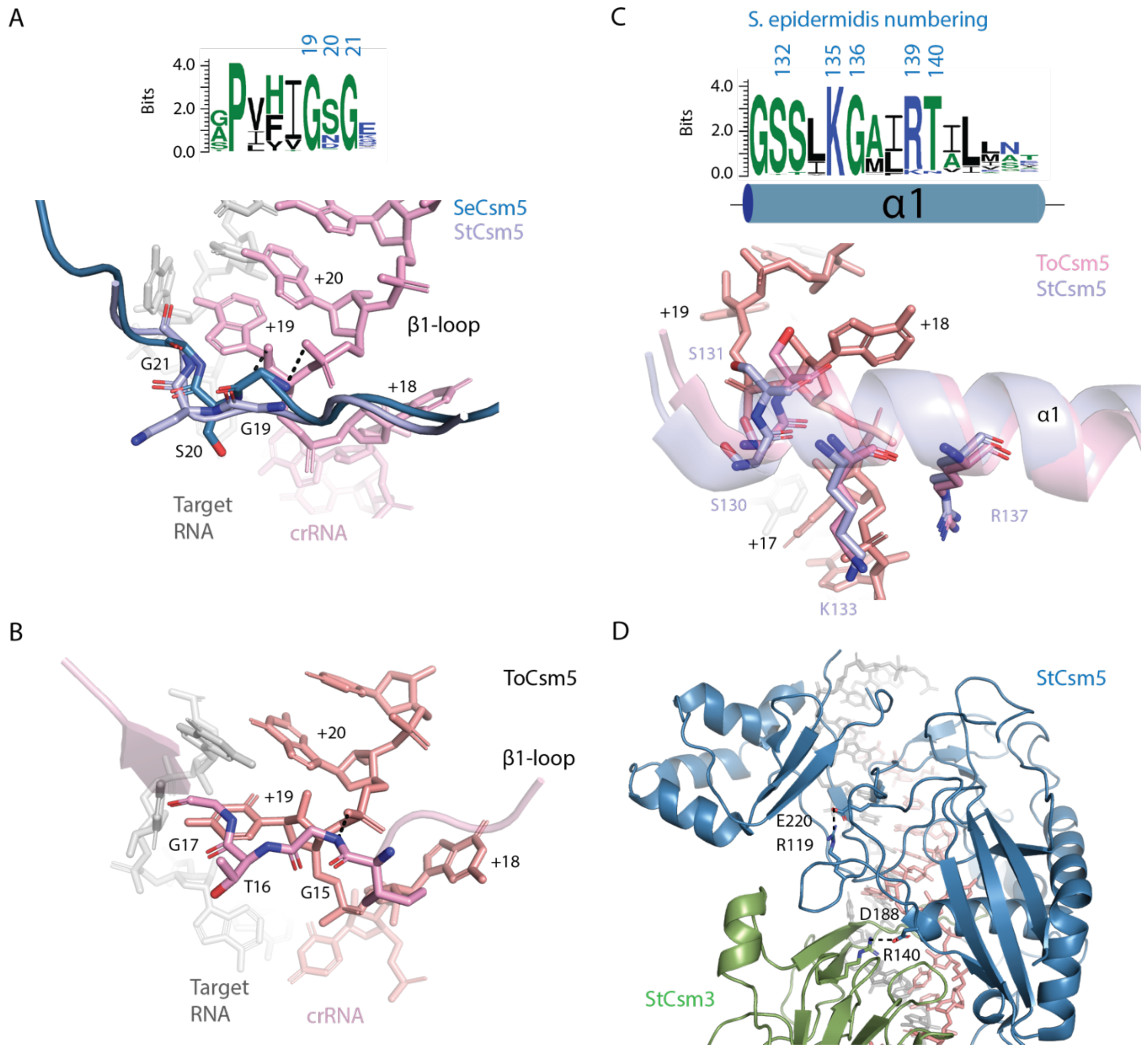


**Figure S9. A comparison of the interactions of bacterial and archaeal Csm5 with target and crRNA.** (*A*) A sharp kink in the peptide backbone of the loop region following β-strand 1 facilitates an interaction with crRNA. A sequence logo depicting the conservation of β1-loop is show above a superposition of *S. epidermidis* Csm5 (SeCsm5) and *S. thermophilus* Csm5 (SeCsm5) highlighting similarities in the interactions of the β1-loop with crRNA. (*B*) The β1-loop of *T. onnurineus* Csm5 (ToCsm5) interacts with crRNA in a conserved manner. (*C*) A sequence logo depicting the conservation of helix-α1 which Csm5 interacts with the flipped +18 nucleotide in a similar manner in bacterial and archaeal Csm5 proteins. (*D*) Csm5 Acidic residues required for crRNA maturation in *S. epidermidis* are conserved in StCsm5 and make similar electrostatic interactions.

**Table S9. SAXS rigid body components**

| Rigid body unit | 6ifu components (chain ID) | rigid body constraints |
|---|---|---|
| 1 | Csm1 (A) K2 – K654, Csm4 (G), Csm3 (F),<br>crRNA (I) A1 – U13, CTR2 RNA (J) A30 – C34 | 4.0 A between unit 1 (Csm1 K654) and unit 2 (Csm1 F655),<br>6.5 A between unit 1 crRNA U13 and unit 2 crRNA C14,<br>6.5 A between target RNA A30 and unit 2 target RNA G29 |
| 2 | Csm1 (A) F655 – K757, Csm3 (E),<br>crRNA (I) C14 – U19, CTR2 RNA (J) A24 – G29 | 6.5 A between unit 2 crRNA U19 and unit 3 crRNA C20,<br>6.5 A between unit 2 target RNA A24 and unit 3 target RNA G23 |
| 3 | Csm2 (C), Csm3 (D),<br>crRNA (I) C20 – U25, CTR2 RNA (J) A18 – G23 | 6.5 A between unit 3 crRNA U25 and unit 4 crRNA C26,<br>6.5 A between unit 3 target RNA A18 and unit 4 target RNA U17 |
| 4 | Csm2 (B), Csm5 (H),<br>crRNA (I) C26 – A34, CTR2 RNA (J) A7 – U17 | |
